# Selective effects of arousal on population coding of natural sounds in auditory cortex

**DOI:** 10.1101/2020.08.31.276584

**Authors:** Charles R. Heller, Zachary P. Schwartz, Daniela Saderi, Stephen V. David

**Affiliations:** Neuroscience Graduate Program, Oregon Health and Science University; Oregon Hearing Research Center, Oregon Health and Science University

## Abstract

The ability to discriminate between complex natural sounds is critical for survival. Changes in arousal and other aspects of behavioral state can impact the accuracy of sensory coding, affecting both the reliability of single neuron responses and the degree of correlated noise between neurons. However, it is unclear how these effects interact to influence coding of diverse natural stimuli. We recorded the spiking activity of neural populations in primary auditory cortex (A1) evoked by a large library of natural sounds while monitoring changes in pupil size as an index of arousal. Heightened arousal increased response magnitude and reduced noise correlations between neurons, improving coding accuracy on average. Rather than suppressing shared noise along all dimensions of neural activity, the change in noise correlations occurred via coherent, low-dimensional modulation of response variability in A1. The modulation targeted a different group of neurons from those undergoing changes in response magnitude. Thus, changes in response magnitude and correlation are mediated by distinct mechanisms. The degree to which these low-dimensional changes were aligned with the high-dimensional natural sound-evoked activity was variable, resulting in stimulus-dependent improvements in coding accuracy.

## Introduction

Humans and other animals are able to discriminate between a multitude of natural sounds. This ability is not static, as the precision of sensory representations by neural activity fluctuates with changes in behavioral state.^1^ Arousal, task engagement, and attention have all been reported to modulate the magnitude and selectivity of single neuron auditory responses,^2–12^ as well as correlated variability across neural populations, often referred to as noise correlations.^13–16^ In general, increased arousal and focused attention are associated with increased response magnitude and decreased noise correlations which are believed to enhance the accuracy of sensory coding.^1, 13, 16, 17^ However, the mechanisms that produce these changes, and the consistency of their effects between different behavioral contexts, are not fully understood.

Recent studies have argued that attention-driven changes in both single neuron responses and correlated activity can be modeled as fluctuations in a single, latent signal that coherently modulates the activity of a subset of neurons. These findings suggest that state-dependent neural population activity occurs in a low-dimensional subspace,^18, 19^ supporting theoretical models in which a single mechanism accounts for changes in single neuron responses and correlated variability.^20, 21^ Fluctuations in arousal, measured by luminance-independent changes in pupil size, modulate neural activity in similar ways to attention,^2, 13, 17^ yet these changes occur independent of attention.^22^ Previous work has not specifically investigated the dimensionality of arousal-dependent signaling and it remains uncertain whether, like other behavioral contexts, it can be explained by a low-dimensional process.

Most studies of population coding accuracy rely on relatively small, simple stimulus sets that drive neural activity in stereotyped ways.^16, 23, 24^ Yet, theoretical work predicts that noise correlations can either enhance or impair coding accuracy, depending on their alignment with the stimulus-evoked activity in the neurons being studied.^20, 25–30^ If the effects of arousal are relatively high-dimensional, meaning that they suppress noise along many different dimensions of neural activity, they should improve coding accuracy of most sensory stimuli equally. Alternatively, if the effects of arousal are confined to a low-dimensional subspace of neural activity, their alignment with sensory-evoked responses should be variable, resulting in stimulus-dependent changes in coding accuracy.

In the present study, we investigated the dimensionality of arousal-dependent signaling and its impact on coding accuracy by recording population activity from primary auditory cortex while presenting a large library of natural sounds. We simultaneously monitored arousal level using pupil size.^2, 31^ To measure population coding accuracy of natural sound stimuli, we developed a novel dimensionality reduction approach.^32, 33^ Overall, arousal improved neural discriminability of natural sounds. However, the degree of improvement varied substantially between stimuli, consistent with the hypothesis that arousal acts on a low-dimensional subspace rather than providing a generalized improvement in coding accuracy. In contrast with attention, modulation of single neuron gain and noise correlations were distinct. These processes operated on different neural populations and timescales. Thus, our results demonstrate that arousal drives robust, selective changes in population coding accuracy across diverse sound stimuli and that these changes act through at least two distinct mechanisms.

## Results

We recorded simultaneous single- and multi-unit activity from primary auditory cortex (A1) using single-shank 64-channel, or dual-shank 128-channel linear silicon probes^34^ (*n* = 371 single-units and *n* = 331 multi-units, Figure 1B). Data were obtained from *n* = 25 recording sites in five awake, head-fixed ferrets. During each recording session, we presented a diverse set of randomly interleaved natural sound excerpts^35^ (*e.g.* Figure 2A) in the acoustic field contralateral to the recording hemisphere (Figure 1A). To monitor spontaneous fluctuations in arousal, pupil size was measured continuously during neural recordings using infrared video^2, 31^ (Figure 1A, B).

**Figure 1.**
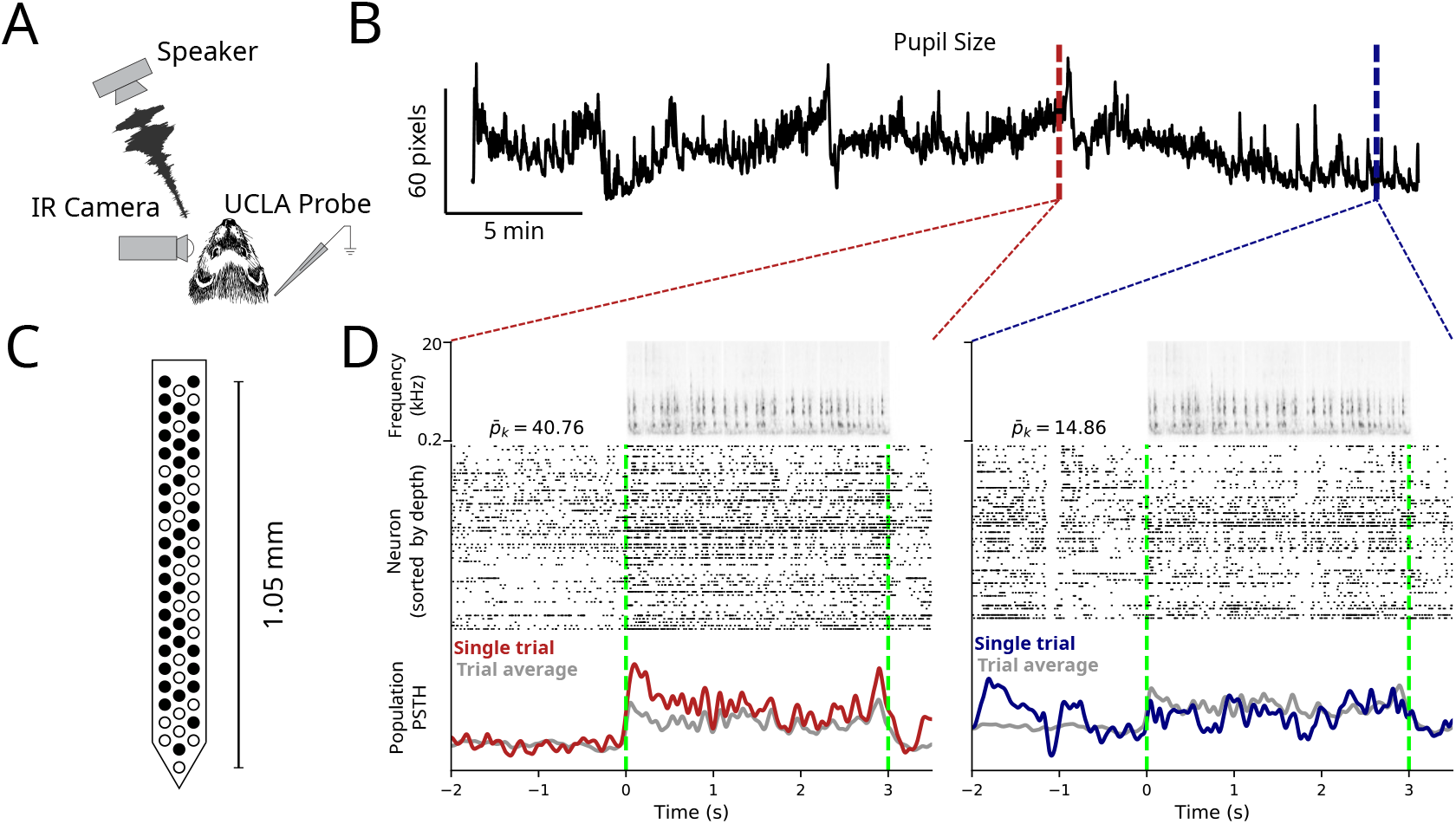
Pupil-indexed arousal modulates neural responses to natural sound stimuli. **A.** Single- and multi-unit activity was recorded from A1 of awake, head-fixed ferrets using laminar electrode arrays during presentation of natural sound stimuli. Pupil size, an index of arousal, was measured simultaneously using infrared video. **B.** Pupil trace from one recording session. Pupil size varied substantially over the course of the recording session, indicating spontaneous transitions between high and low arousal states. **C.** Schematic of 64-channel laminar probe used to record neural activity. Filled circles represent electrode channels on which at least one unit was detected during the same session (*n* = 55 total units). **D.** Top: Spectrogram of one 3 s natural sound excerpt which was presented multiple times during the recording session. Middle: Population raster plot of spiking activity by all simultaneously recorded units during a single stimulus presentation when pupil was large (left, red arrow in B) and when pupil was small (right, blue arrow in B). Bottom: Population peri-stimulus time histogram (PSTH) response, averaged across units during the single trial (red / blue) indicated in B and averaged over all repetitions of this stimulus (gray). 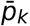 indicates mean pupil size on each respective trial, *k*.

**Figure 2.**
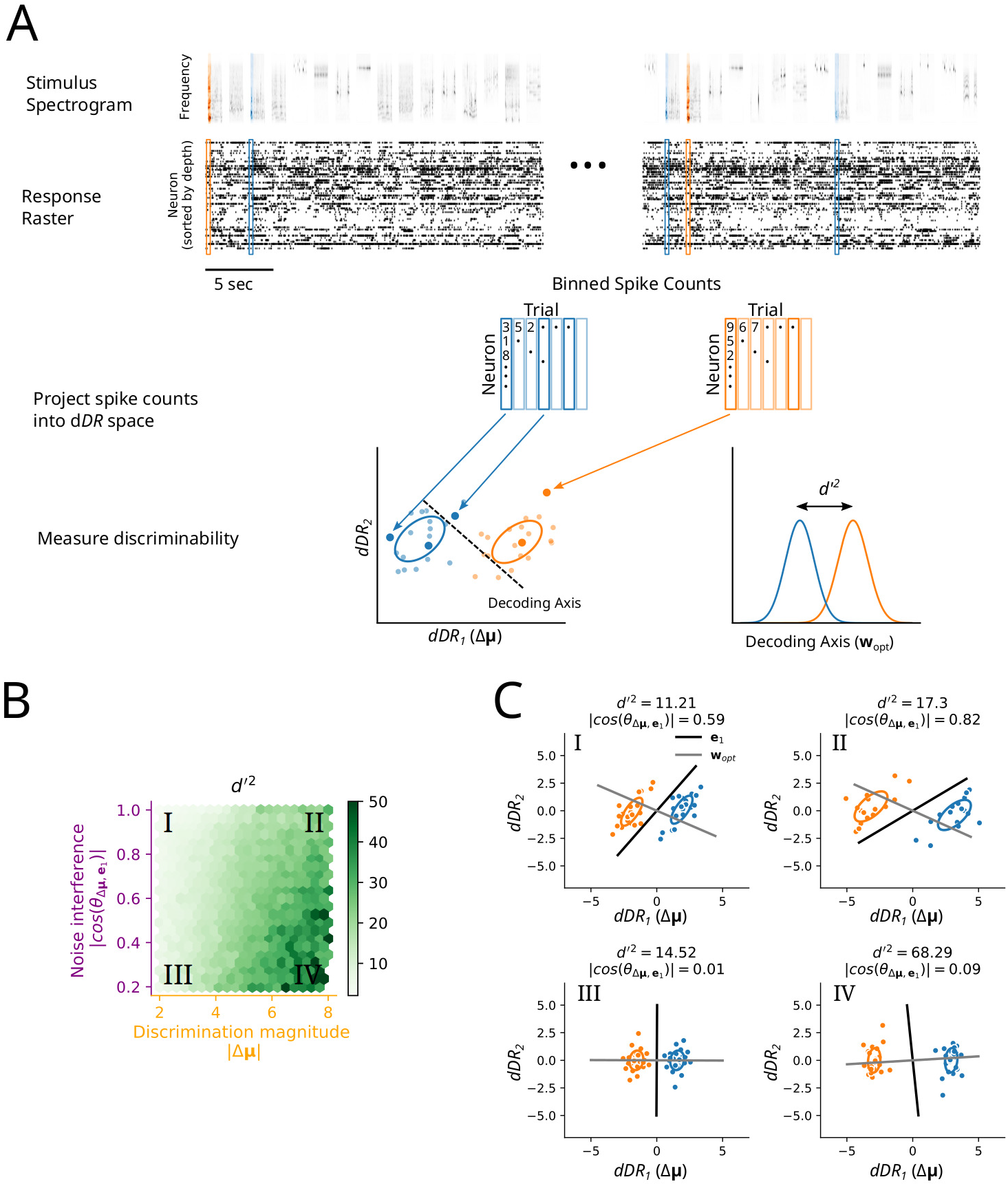
Natural stimulus discriminability in A1 populations varies smoothly across the sensory space. **A.** Procedure for measuring discriminability between natural sound pairs. Spectrogram and response raster from one example recording session are shown for exemplar early (left) and late (right) trials in the experiment. For each 250 ms segment of sound stimuli, spike counts in the corresponding time window were binned and counted as shown. For each pair of unique sound segments (blue vs. orange), the spike counts were then projected into *dDR* space, where *d*′^2^ was measured. **B.** Heat map indicates neural discriminability, *d*′^2^, averaged across stimulus pairs and recording sites. Values were binned according to discrimination magnitude (*x*-axis) and noise interference (*y*-axis). **C** I-IV. Single-trial responses to four example sound pairs. Each dot represents the weighted sum of neural population activity on a single trial. Ellipses indicate standard deviation across trials. Example stimulus pairs shown were all collected from the same neural population during a single recording session. Each panel shows a pair of stimuli from a different quadrant (I - IV) in B. **e**_1_ (black line) represents the first eigenvector of the noise in the *dDR* space and **w**_*opt*_ (grey line) represents the optimal decoding axis for separating the two stimulus classes (blue vs. orange).

In ferret A1, changes in pupil size are associated with mostly monotonic changes in neural firing rate.^2^ Therefore, to assess overall pupil-related changes in activity, we split the neural data in half based on the median pupil size across each experiment (large pupil/high arousal vs. small pupil/low arousal). Transitions between high and low pupil-indexed arousal were accompanied by changes in the excitability of individual neurons and in the strength of correlations between neurons (Figure 1D). When pupil was large, responses to the same sound were stronger and more reliable than when pupil was small. Across all recorded units, both the baseline firing rate and evoked response gain were positively associated with pupil size (Figure S1). Additionally, during large pupil trials population activity was desynchronized relative to the low arousal state; coordinated, stimulus-independent fluctuations in the population PSTH were primarily observed during small pupil trials only. In support of this observation, we found that pairwise noise correlations were significantly reduced in the high arousal state (*ρ_small_* = 0.047 *±* 0.005 vs. *ρ _large_* = 0.043 *±* 0.005, *p* = 0.035, Bootstrap test, *n* = 25 recording sessions, 318 *±* 94 unit pairs per session; Figure S1). These results are consistent with previous reports in ferret,^2^ mouse,^13, 17, 31^ and primate.^36^

### The impact of correlated variability on population coding accuracy varies across sensory stimuli

We directly measured the impact of noise correlations on coding of natural sounds using *d*′^2^, a well-established metric of neural population discriminability.^26, 27, 32, 37–39^ *d*′^2^ describes the ability of population activity to discriminate between two stimuli using an optimal linear decoder. Its value depends both on the mean, stimulus evoked activity and on the variability of responses across trials, or noise correlations, that project along the optimal decoding axis.^26–28^

To prevent overfitting to noise in the high-dimensional population data despite having relatively few repetitions of each stimulus, we performed dimensionality reduction.^32, 33^ Before measuring *d*′^2^, we projected neural responses onto a two-dimensional plane in neural state space. This projection, which we call decoding-based Dimensionality Reduction (*dDR*, Figure S3), was defined by two axes: the sensory discrimination axis (*dDR*_1_ / Δ***μ***) and the global noise axis (*i.e.* the first principal component of noise correlations, see Methods). In addition to preventing overfitting to single trial noise in *d*′^2^ estimates (Figure S4), *dDR* facilitated visualization of the high dimensional neural population data (Figure 2A, C). Increasing the dimensionality of the *dDR* space beyond two-dimensions by including additional noise components only increased cross-validated estimates of *d*′^2^ marginally and did not impact the effect of arousal on *d*′^2^, which is the main focus of this work (Figure S6).

Neural discriminability was strongly stimulus-dependent (Figure 2B, C). To determine the source of stimulus dependence, we characterized each stimulus pair by two metrics: discrimination axis magnitude and noise interference (Figure 2B). Discrimination axis magnitude was defined as the vector magnitude of Δ***μ***, which described the amount of sensory information contained in the trial-averaged activity. Noise interference was defined as the cosine similarity between Δ***μ*** and the correlated variability axis (**e**_1_, i.e., the first principal component of stimulus-independent activity in *dDR* space). Thus, noise interference was a stimulus pair specific metric that described the extent to which single trial variability interfered with the readout of sensory information. These two metrics are used throughout the remainder of this work. To reference their definitions, see Methods: Glossary.

Across the full set of stimulus pairs, we observed substantial variability in both discrimination magnitude and noise interference (Figure S5), which in turn led to changes in discrimibaility (Figure 2B). For pairs of stimuli with large discrimination magnitude and low noise interference, *d*′^2^ was large (Figure 2C.IV) and for pairs with small discrimination magnitude and high noise interference, *d*′^2^ was small (Fig 2B, C.I). To quantify this, we regressed *d*′^2^ against the per-stimulus noise interference and discrimination axis magnitude (Figure 3E, Eqn. 10). For all sites, noise interference coefficients were negative and discrimination magnitude coefficients were positive (*β_noise_* = *−* 0.39 *±* 0.03*, p* = 0.000071*, U* = *−*3.97*, n* = 11 recording sessions, Mann-Whitney U Test; *β_discrimination_* = 0.80 *±* 0.02*, p* = 0.000071*, U* = 3.97*, n* = 11 recording sessions, Mann-Whitney U Test), indicating that this pattern was consistent across experiments.

Our results illustrate that baseline neural discriminability of natural sounds varies systematically across the stimulus space. The variation of *d*′^2^ with respect to noise interference emphasizes that correlated variability has very little impact on discrimination for stimulus pairs, but is critically important for others. This dependence of discriminability on noise interference persisted after controlling for discrimination magnitude, demonstrating that the effect of noise correlations varies even across pairs of stimuli that differ similarly in their mean response.

**Figure 3.**
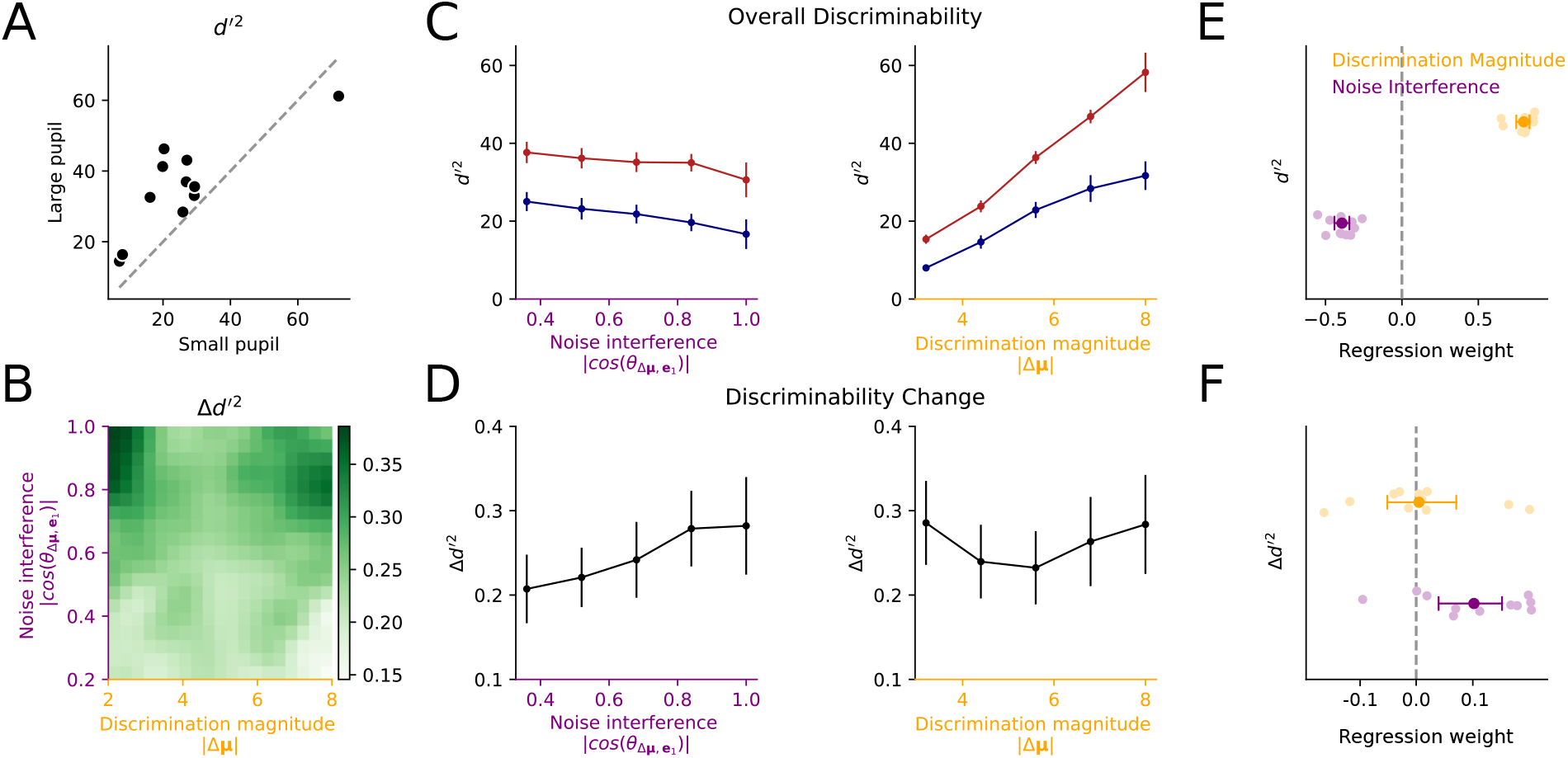
Arousal improves discrimination of natural sounds. **A.** Scatter plot compares mean discriminability (*d*′^2^) across natural sound pairs for small (low arousal) versus large (high arousal) pupil trials. Each point indicates the mean across all stimulus pairs presented during a single recording session. Mean discriminability was nearly always greater during large pupil trials (*p* = 0.02*, W* = 7*, n* = 11 sessions, Wilcoxon signed-rank test). **B.** Heatmap shows the relative change in discriminability for large vs. small pupil (Δ*d*′^2^) as a function of discrimination axis magnitude (|Δ***μ***|) and noise interference (|*cos*(*θ*_Δ***μ***,**e**1_)|). Results are smoothed by a Gaussian filter of width two bins. **C.** Average *d*′^2^ for large and small pupil trials grouped into five evenly spaced bins by noise interference (left) or discrimination magnitude (right). Points / error bars indicate the mean / standard error across recording sessions. **D.** Arousal-related fraction improvement in discriminability (Δ*d*′^2^) for each session, plotted as in C. **E** Regression coefficients computed per recording session for dependence of *d*′^2^ on discrimination magnitude and noise interference. Error bars show bootstrapped 95% confidence interval across sessions. *β_Noise_* = −0.39 ± 0.03*, p* = 0.000071*, U* = −3.97*, β_Discrimination_* = 0.80 ± 0.02*, p* = 0.000071*, U* = 3.97*, n* = 11 recording sessions, Mann-Whitney U test. **F.** Coefficients for pupil-dependent changes in discriminability, m*d*′^2^, plotted as in (E). *β_Noise_* = 0.102 ± 0.029*, p* = 0.001*, U* = 3.25*, β_Discrimination_* = 0.004 ± 0.030*, p* = 0.718*, U* = 0.36*, n* = 11 sessions, Mann-Whitney U test.

### Arousal selectively improves neural discrimination of natural sound stimuli

We used the *dDR*-based decoder to determine how pupil-indexed arousal modulates population coding accuracy in A1. For each pair of stimuli, we computed a single decoding axis across all pupil states, as above (**w**_*opt*_, Eqn. 9). We then split the data from each stimulus pair in half based on the median value of pupil across sound repetitions and computed *d*′^2^ separately for small and large pupil conditions. The use of a single decoding axis, combined with the median split of data based on pupil size, prevented biasing *d*′^2^ for one or the other pupil condition. Across stimulus pairs, we found that discriminability of natural sounds was improved on average in large pupil, high arousal states (Figure 3A). This change was significant across the population (*p* = 0.02*, W* = 7*, n* = 11 recording sites, Wilcoxon signed-rank test). Moreover, the magnitude of the effect was correlated with the amount by which pupil varied within an experiment, consistent with it being driven by pupil-indexed fluctuations in arousal (Figure S7, *r* = 0.63*, p* = 0.029*, n* = 11 recording sessions, permutation test).

Our analysis of overall discriminability revealed substantial variation in the impact of noise correlations on *d*′^2^ across stimulus pairs (Figure 2B). Therefore, we next considered the possibility that pupil-dependent changes in coding accuracy also depend on these same features. To test this, we computed the fraction change in discriminability between large and small pupil conditions, m*d*′^2^ (Eqn. 12), which ranged from ***−***1 to 1 and could be directly compared across stimulus pairs and recording sites. We binned stimulus pairs according to the discrimination axis magnitude and noise interference (as in Figure 2B) and averaged m*d*′^2^ within bins and across recording sites (Figure 3B). Unlike overall discriminability, improvements did not depend on discrimination magnitude (Figure 3D, right). However, m*d*′^2^ was positively correlated with noise interference; for stimulus pairs with high noise interference, increased arousal led to a greater relative improvement in discriminability (Figure 3D, left). We quantified these axis-dependent changes in m*d*′^2^ using linear regression. m*d*′^2^ and noise interference were positively correlated in 8*/* 11 recording sites (*p <* 0.05, t-test), and this correlation was significant across experiments (*β_noise_* = 0.102 *±* 0.028*, p* = 0.001*, U* = 3.25*, n* = 11 recording sessions Mann-Whitney U Test; Figure 3F). No consistent relationship was observed for discrimination magnitude (*β_discrimin tion_* = 0.004 *±* 0.030*, p* = 0.718*, U* = 0.36*, n* = 11 recording sessions, Mann-Whitney U Test; Figure 3F). These results demonstrate that improvements in stimulus discriminability associated with increased arousal are largest for stimulus pairs where correlated variability aligns with the discrimination axis. The selectivity of this effect suggests that state-dependent changes are low-dimensional and that they align with the most prominent axis of correlated variability.

### Arousal improves discriminability through a combination of increased response gain and reduced correlated variability

Changes in either single neuron response magnitude or noise correlations (Figure 1-S1) could lead to enhanced discriminability in the large pupil condition. We reasoned that three population level factors could capture these effects: discrimination axis (signal) magnitude, shared noise variance, and noise interference (Figure 4A). Discrimination axis magnitude is directly related to *d*′^2^ (Figures 2B and 3C) and is measured using trial-averaged activity. Thus, pupil-dependent changes in this value only reflect changes in the evoked response magnitude of single neurons. Shared noise variance, on the other hand, is independent of trial-averaged responses and reflects the strength of trial to trial variability. Noise interference, as we discuss above, reflects the interaction between these two quantities.

**Figure 4.**
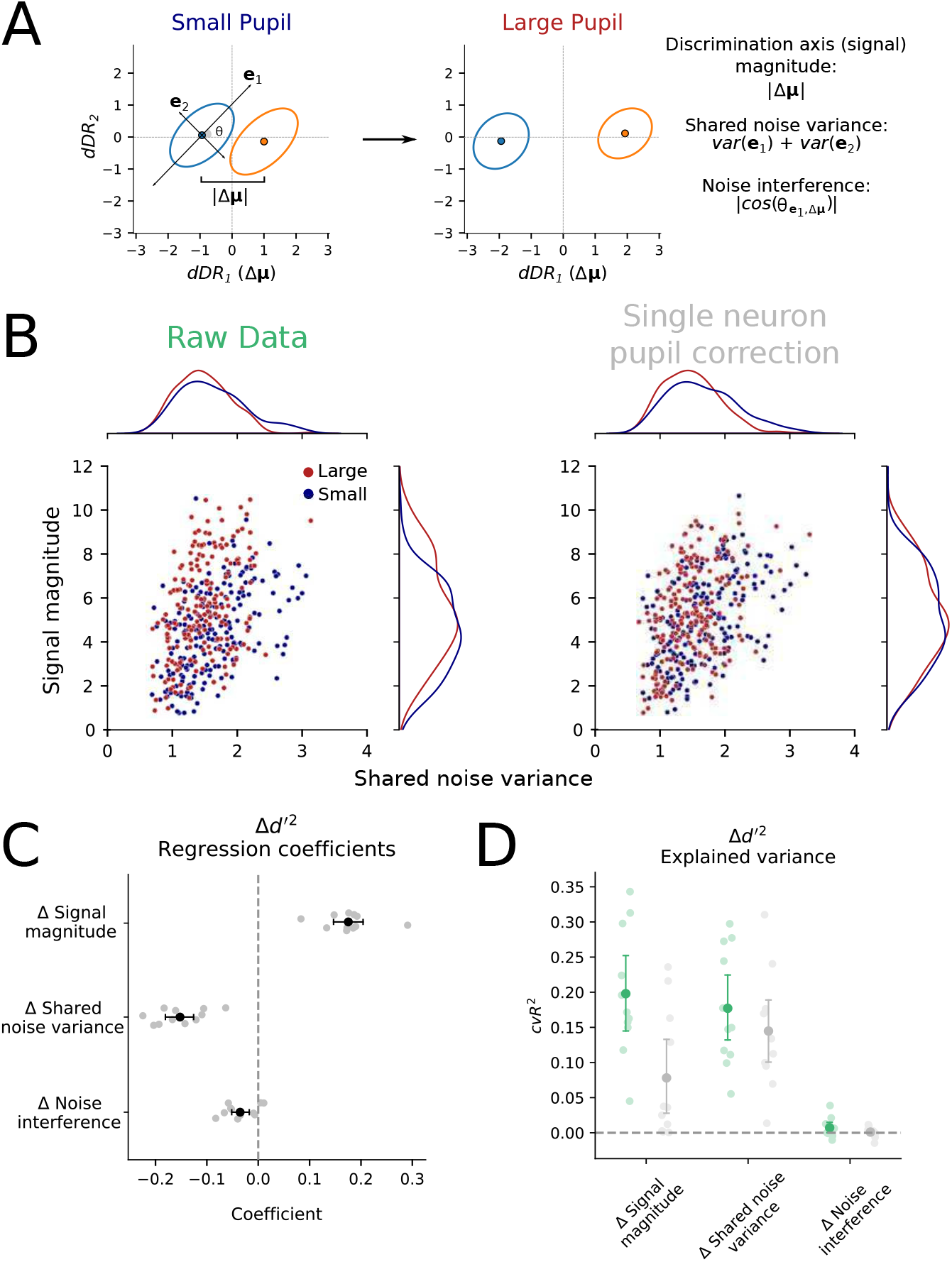
Arousal-dependent changes in both mean activity and trial to trial variability impact stimulus discriminability. **A.** Cartoon schematic of typical arousaldependent modulation of population responses to two stimuli (blue and orange) in *dDR* space. A change in mean response affects the distance between ellipsis centers, and a change in correlated variability affects their size. **B.** Scatter plot of signal magnitude (mean difference along discrimination axis) versus shared noise variance for each stimulus pair. Color indicates small versus large pupil conditions, randomly sub-sampled to facilitate visualization (*n* = 200*/* 17, 778 stimulus pairs). **C.** Linear regression was used to model changes in discriminability (*d*′^2^) as a function of changes in signal magnitude, shared noise variance, and noise interference (Eqn. 13). Each gray point represents the regression coefficient for data from one experiment (*n* = 11). Error bars indicate standard error of the mean coefficient across experiments. **D.** Cross-validated change in *d*′^2^ explained by each regressor (*cvR*^2^, Methods) before (green / raw) and after (grey / pupil correction) correcting for pupil-dependent changes in single neuron evoked responses. Error bars represent standard error of the mean *cvR*^2^ across experiments.

Inspection of changes in these factors for individual stimulus pairs showed that discrimination magnitude increased in the large pupil condition and shared noise variance decreased (Figure 4B, left). We used a regression model to determine the extent to which each could explain changes in *d*′^2^ (Figure 4C, D). Consistent with a contribution to the change in discriminability, a positive change in discrimination magnitude and a decrease in noise variance both predicted an increase in *d*′^2^ (Figure 4C). Each of these factors contributed significantly to the improved discriminability during high arousal conditions (Figure 4D, discrimination (signal) magnitude: mean cross-validated *R*^2^ = 0.198*, p* = 0.000071*, U* = 3.97, shared noise variance: mean *R*^2^ = 0.177*, p* = 0.000071*, U* = 3.97*, n* = 11 recording sessions, Mann-Whitney U test). Noise interference, however, changed only marginally between arousal states and did not explain substantial variability in *d*′^2^ (mean cross-validated *R*^2^ = 0.007*, p* = 0.28*, U* = 1.08*, n* = 11 recording sessions, Mann-Whitney U test).

Both independent, single neuron variance, as well as noise correlations, could impact discriminability. As validation that the our measurement of shared noise variance corresponded primarily to modulation of noise correlations, and not changes in single neuron variance, we simulated population activity. In the simulation, the mean and variance of single neurons was matched to the values measured for the small and large pupil states, but noise correlations were held fixed (Figure S9, Methods). These simulated data were unable to account for the actual observed changes in *d*′^2^, while a simulation that also incorporated changes in noise correlations accurately reproduced the raw data. Thus, a combination of changes in the trial-averaged, evoked activity of single neurons and in correlated variability between neurons is needed to explain decoding improvements.

### Arousal-dependent changes in gain and correlated variability operate on distinct timescales and neural populations

Studies of selective attention have shown that attention-related changes in response gain of single neurons and the strength of correlated variability are related. In this framework, changes in correlated variability reflect changes in the variance of coherent modulation of the sensory gain of single neurons across the population.^18, 19^ Thus, a single, shared mechanism appears to mediate changes in both gain and correlation. We wondered if a single mechanism also produced effects of arousal or if changes in gain and correlated variability were mediated by separate mechanisms.

If arousal impacts single neuron activity and correlated variability via a single mechanism, we reasoned that removing all pupil-dependent changes in single neuron spiking activity should also remove population-level changes in both discrimination axis magnitude and shared noise variance (Figure 4B). To test this hypothesis, we generated a pupil-corrected data set in which all pupil-explainable response variability for each unit was subtracted from its raw response (Eqn. 8). We then measured the pupil-dependent discrimination axis magnitude, shared noise variance, and stimulus discriminability for the pupil-corrected data. The single neuron correction abolished changes in discrimination axis magnitude, but changes in shared noise variance were largely unaffected (Figure 4B). These results argue that arousal does not modulate correlated variability by reducing shared fluctuations in arousal-dependent gain.^18, 19^ Instead, increased arousal suppresses an independent source of correlated variability. These separable mechanisms are distinct from reports for selective attention, where changes in response gain and correlated variability are thought to be produced by a single shared modulator.

To further distinguish mechanisms that impact discrimination magnitude and correlated variability, we used a bandpass filter to partition the data into distinct temporal frequency bands and measured noise correlations separately in each band (Figure 5A, B). Breaking the data into frequency bands allowed us to identify the timescale over which the correlation effects operate. Noise correlations were largest overall in low-frequency bands (< 0.5 Hz), reflecting slow, coordinated fluctuations in the population activity.^40^ Pupil-indexed fluctuations in arousal occur at this slow timescale. Thus, we expected the effects of coherent single neuron modulation to be largest in this band. Indeed, the pupil correction reduced these slow noise correlations significantly (*p* = 0.00003*, W* = 6*, n* = 25 recording sessions, Wilcoxon signed-rank test, Figure 5A) but not in the higher frequency bands, *>* 2 Hz. In contrast, arousal-dependent changes in correlation magnitude were restricted to the higher frequency bands, ranging between 0.5 and 25 Hz (Figure 5B, *p* = 0.019, 0.002, 0.007, 0.00005*, W* = 76, 51, 63, 11*, n* = 25 recording sessions, Wilcoxon signed-rank test). Therefore, we conclude that the arousal-dependent changes in correlated variability are due to modulation of a process that operates on a faster timescale, distinct from the change in arousal itself.

**Figure 5.**
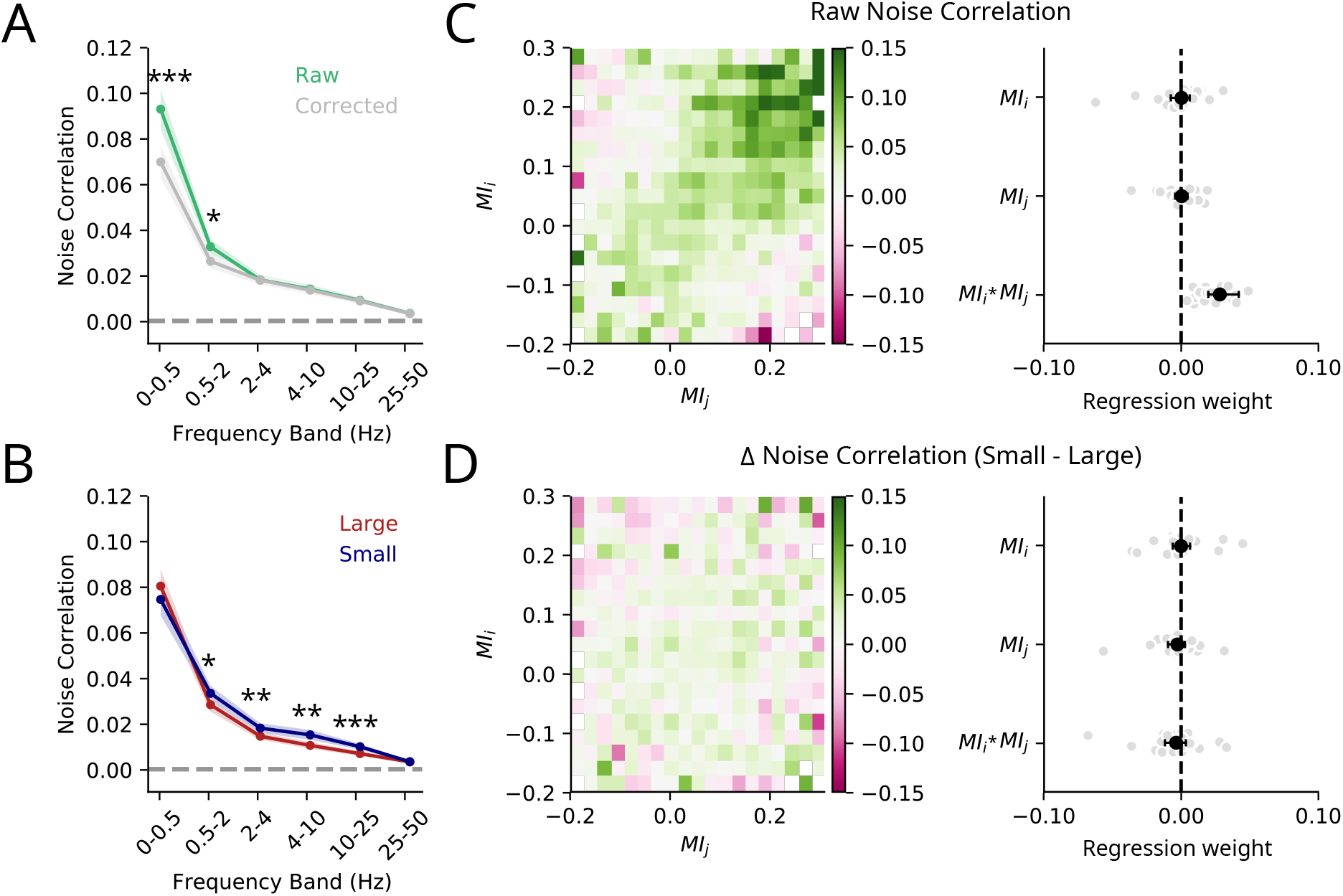
Arousal-dependent changes in evoked response magnitude and correlated variability operate on distinct timescales and neural populations. **A.** Noise correlations, computed with and without pupil correction, split into six non-overlapping temporal frequency bands. To minimize bias from one recording session, noise correlations were averaged across neuron pairs within session before measuring significance of the difference between conditions (\*\*\**p <* 0.001, \*\**p <* 0.01, \**p <* 0.05, Wilcoxon signed-rank test, *n* = 25 recording sessions). Shading indicates standard error across recording sessions. **B.** Pupil-corrected noise correlations measured after splitting data by median pupil size, plotted as in A. **C.** Left: Heatmap of mean noise correlations (using 4Hz temporal bins, as in Figure S1), grouped into 20 evenly spaced *MI* bins for each pair of units (*i*, *j*). Right: Regression coefficients, predicting overall noise correlations from *MI* of individual neurons (*MI_i_* and *MI_j_*) and their interaction (*MI_i_* ∗ *M_j_*). Gray dots represent single recording sessions; error bars represent 95% confidence intervals across sessions. **D.** Same as C but for change in noise correlations (m noise correlations) between small and large pupil.

If changes in correlated variability are in fact distinct from single neuron effects, the two processes may also operate on different subpopulations of neurons. To determine if this was the case, we compared the magnitude of pupil-dependent modulation observed in each cell with the degree to which arousal state impacted noise correlations for that cell. Single cell effects were quantified by the pupil modulation index (MI) (Eqn. 4). A value of MI= 1 indicated a unit that only responded when pupil was large and MI= 0 indicated no change in responsiveness between large and small pupil. While MI was able to predict noise correlations between two units, it contained no information about the arousal-dependent change in noise correlations (Figure 5C, D). Thus, in addition to acting on distinct timescales, each process also operates on unique populations of neurons in A1, confirming that single neuron changes are distinct from changes in correlated variability.

### Arousal-dependent reduction of correlated variability is low dimensional

Having determined that arousal-dependent changes in correlated variability operate through a distinct mechanisms from changes in individual neuron excitability, we finally sought to determine the dimensionality of the mechanism producing the correlations. Specifically, we determined if changes in noise correlations could be described by a low dimensional latent process, or if they reflected more complicated modulation of high-dimensional neural coupling across the population. For each session, we computed the axis of maximal change in noise correlations between large and small pupil (Figure 6A-D). In most cases, this pupil modulation space was low-dimensional; in 9*/* 11 recording sessions, we found only one significant dimension along which noise correlations changed between large and small pupil (*p <* 0.05, permutation test, Methods). This dimension was consistently aligned with the first principal component of the pooled noise data (6E, F), suggesting that arousal modulates correlated variability along, low-dimensional, high variance dimensions in the data.

**Figure 6.**
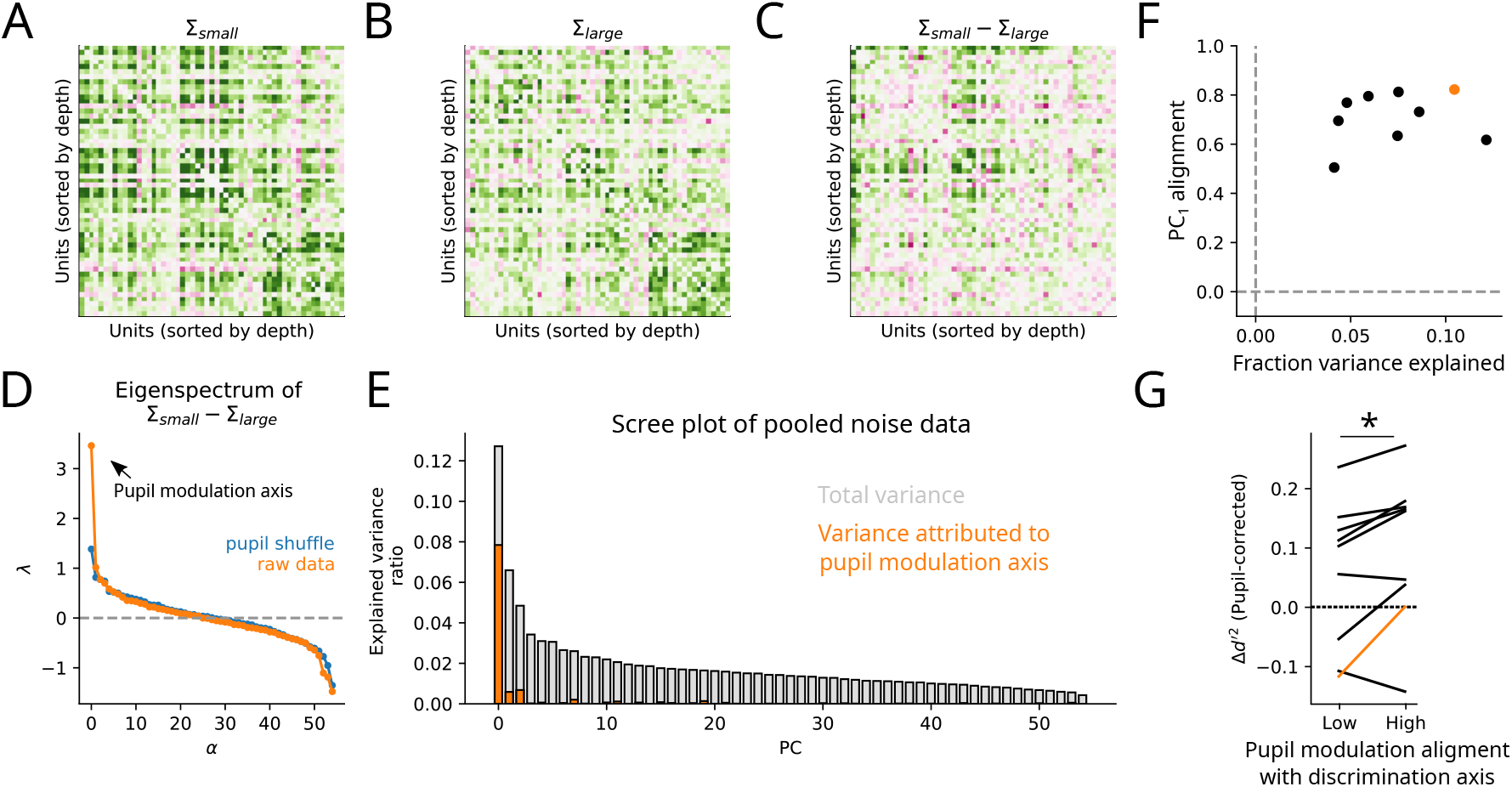
Pupil-related changes in noise correlations are low-dimensional and predict changes in stimulus discriminability. **A.** Matrix of pairwise noise correlations between all units during the small pupil condition in one recording session. **B.** Same as in A for large pupil condition. **C.** Change in noise correlations, computed as the difference between matrices in A and B. Color scale shared between panels A-C. **D.** Eigenvalues of the difference matrix in C (orange). The first eigenvalue corresponds to the pupil modulation axis. Blue line shows the noise floor, computed by shuffling pupil size before classifying large vs. small pupil conditions. Only sites with at least one significant eigenvalue were included subsequently in panels F/G (9/11 sites, *p <* 0.05, permutation test). **E.** Scree plot shows variance explained ratio for each principal component. Orange shading indicates the portion of variance along each PC that can be attributed to the pupil modulation axis. The large orange component in the first position indicates that the pupil modulation axis largely aligns with the first principal component. **F.** Scatter plot compares fraction of total variance in the population activity explained by the modulation axis (*x*-axis) against the ratio of modulation axis variance to *PC*_1_ variance (*y*-axis), *i.e.*, the ratio between heights of the orange and grey bars for *PC*_1_ in panel E. Example site from A-E is shown in orange. **G.** Change in *d*′^2^ between pupil conditions after performing single neuron pupil correction for each site. Mean change is computed across stimulus pairs after first grouping each pair by the alignment of the pupil modulation axis with its discrimination axis (median split between low and high alignment). Stimulus pairs with high alignment (measured with cosine similarity) showed greater discrimination improvements (μ_*high*_ = 0.099 *±* 0.042, μ_*low*_ = 0.057 *±* 0.041*, p* = 0.028*, n* = 9, Wilcoxon signed-rank test).

Given that arousal-dependent changes are usually restricted to a single dimension, we hypothesized that their impact on discriminability will depend on how well that axis aligns with the sensory discrimination axis. To test this, we compared the pupil modulation axis to the sensory discrimination axis (Δ***μ***) for each stimulus pair. Indeed, this was the case. For pairs of stimuli where the cosine similarity between Δ***μ*** and pupil modulation axis was high, we observed that the effect of correlated variability changes on *d*′^2^ were larger (Figure 6G, *p* = 0.028*, W* = 4*, n* = 9, Wilcoxon signed-rank test).

Based on these results, we conclude that arousal-dependent changes in correlated variability are restricted to a relatively low-dimensional subspace. These changes increase sound discriminability when it is aligned with the discrimination axis, a scenario that only occurs for a subset of sensory discriminations.

## Discussion

Previous studies have suggested that behavior-dependent modulation of neural population coding operates in a low-dimensional space.^18, 19, 21, 22, 40, 41^ That is, signals reflecting behavioral state are well-described by processes that modulate the activity of many neurons coherently and thus produce correlated variability in sensory responses. However, most previous work has utilized relatively small, focused stimulus sets. This raises questions about whether the observed low-dimensional processes are a consequence of the stimuli tested, or if they are a general feature of state-dependent modulation. These questions are critical for understanding population coding of sensory stimuli. Theoretical studies have long shown that correlated variability can impact coding accuracy, but only if it aligns with the sensory tuning of neurons in the population.^27^ Thus, the dimensionality of the mechanisms driving correlated variability and how they interact with sensory selectivity is critical for understanding their impact on sensory processing.

In the case of pupil-indexed arousal, we found that correlated activity is modulated in a low-dimensional subspace of primary auditory cortex (A1) which we found to be distinct from the arousal-dependent changes in single neuron responses. These results were consistent across a diverse set of natural sound stimuli. The effect of arousal on neural discrimination of sounds varied substantially with the sound stimulus, as predicted for a low-dimensional signal interacting with high-dimensional stimulus-evoked activity.

### Effects of shared intrinsic variability on discriminability are stimulusdependent

Correlated, intrinsic variability within neural populations is ubiquitous in cortex. Even before this phenomenon was observed experimentally, substantial efforts were made to develop a theoretical understanding of how correlated activity might affect coding by neural populations.^26, 37, 42–47^ This early work established that correlated variability can interfere with the brain’s ability to accurately discriminate sensory stimuli. Therefore, experimental characterization of this phenomenon is critical to fully understand neural population codes.

Although evidence for intrinsic correlation is widespread, experimental studies have provided conflicting evidence as to whether or not it does in fact interfere with population coding.^20, 25, 28–30, 32, 38, 39^ There are at least two reasons why the reported effects of correlated variability might vary across studies. First, in some cases activity along dimensions containing interfering noise could have very low variance. In this case, measuring the noise reliably would require recording large amounts of data, both over many neurons and over many trials, a methodology that has only recently become feasible.^32, 38, 39^ A second possibility is related to the fact that the impact of correlated noise depends on the tuning of neurons in the population being read out and its relationship with the noise space.^26, 28, 30^ In this case, discrepancies in previous work might be explained by differences in the neural populations that were sampled or in the stimuli that were tested. Because the effects of intrinsic noise may depend on the stimuli that are presented, it is important to characterize coding accuracy across diverse sets of stimuli. Indeed, our results showed that the effects of correlated variability on coding are highly dependent on the stimuli tested.

Because there is a trade-off between the number of stimuli that can be presented and the number of times that each can be repeated during a single recording session, questions about stimulus-dependent changes in population coding are difficult to completely address in a single study. Unlike recent work,^32, 38, 39^ we measured neural responses to a large set of stimuli over a relatively small number of repeats and neurons. Thus, we could not measure low-variance dimensions and draw strict conclusions about the presence (or absence) of information-limiting noise.^26^ Instead, by developing a novel dimensionality reduction approach (*dDR*), we were able to reliably estimate the interaction between the dominant, high-variance noise dimension and sensory discrimination across a large acoustic stimulus space. This approach revealed substantial variability of arousal dependent changes in coding within each recording site. This highlights the practical benefit of dimensionality reduction techniques for studying neural population dynamics across a diversity of stimulus and behavioral contexts.^48^

### State-dependent coding in auditory cortex

It is increasingly clear that neural activity in primary sensory regions of the brain is significantly modulated by non-sensory variables, including arousal.^2, 13, 17, 40, 49^ Arousal here refers to spontaneous changes in alertness as measured by pupil diameter, even in the absence of a behavioral task.^49^ Similar to previous work, we find that increased arousal is associated with enhanced excitability and reduced noise correlations in A1.^2, 13, 31^ These effects boost the neural signal to noise ratio in V1^17^ and improve population coding accuracy of tonal stimuli in A1.^13^

Building on this previous work, we explored the effects of arousal on population coding accuracy across a large space of natural sounds. Consistent with prior results, increases in pupil-indexed arousal led to improved discriminability between sounds on average. However, the relative magnitude of this improvement varied substantially across stimuli. Improvements were largest when correlated variability interfered with sensory discrimination. Thus, a suppression of correlated variability and increase in evoked response rates during high arousal states may gate behavioral states that require improved perceptual discrimination, like selective attention.

This strong stimulus dependence highlights the importance of a systematic exploration of state-dependent changes in neural coding across the sensory response space. Parametric stimuli might be used to more systematically probe sound feature representations across a range of behaviorally relevant stimuli. For example, one study of auditory processing began to address this question in anesthetized animals.^23^ In this work, Kobak et al. measured population coding in A1 of sounds that varied along two dimensions: Inter-aural level (ILD) and absolute binaural level (ABL). By inducing different states of cortical activation with urethane anaesthesia, the authors demonstrated that in the awake (desynchronized) state, noise and signal subspaces shift to become orthogonal, thereby facilitating accurate encoding across both ILD and ABL. Extending this approach to spectro-temporally varying and behaviorally relevant naturalistic stimuli^50^ will be critical for a complete understanding of state-dependent population coding.

### Separate mechanisms drive arousal-dependent changes in single neurons and correlated neural variability

Recent studies of selective attention have suggested that correlated variability results from the coherent modulation of many neurons by an intrinsic behavioral state variable.^18–20^ In a recent study by Denfield et al., macaques were trained on a visual change-detection task in which the stability of spatial selective attention was manipulated between behavioral blocks.^19^ Because the gain of evoked responses in visual cortex is known to be modulated by attention,^51^ the authors proposed that the magnitude of correlated variability should be highest when attention itself was most variable, as changes in gain are shared across neurons within the receptive field. Indeed, when animals were required to switch attention between multiple locations within a behavioral block, noise correlations were strongest. This idea of correlations produced by a shifting spotlight of attention is consistent with previous characterizations of neural population activity and attention^18^ and agrees well with theoretical work.^20, 21^ These findings offer a parsimonious explanation for why gain changes are accompanied by a reduction in noise correlations during traditional cued change-detection tasks, where attention is focused stably on a single spatial location.^16^

Unlike the case of visual selective attention, we found that arousal-dependent modulation of evoked rates and noise correlations in auditory cortex could be fully dissociated, and thus did not arise from a common signal. Modulation of single neuron activity was slow, on the order of many seconds. These slow changes may be analogous to previously described drift signals in cortex.^18, 22^ Correlated variability between neurons, however, also operated on a faster timescale, from tens to hundreds of milliseconds. The magnitude of these faster correlations was modulated by the slow arousal signal, but we could not predict changes in correlated variability from modulation of single neurons. Instead, neurons undergoing slow changes in excitability and those undergoing modulation of faster noise correlations comprised two independent subpopulations.

Although it was not feasible to directly isolate the circuitry underlying these distinct effects in the current study, we propose that they may arise through a combination of neuromodulation and intracortical feedback. Several studies have shown a strong correlation between slow fluctuations in pupil diameter and brain-wide release of norepinephrine and acetlycholine,^52^ making them good candidates for mediating the slow changes in response baseline and gain across individual neurons. The decrease in correlated activity, on the other hand, may arise due to modulation of feedback from other cortical areas that are themselves targeted by the same neuromodulatory signals.

Intracortical pathways to auditory cortex have been identified from multiple areas, including visual,^53^ motor^54^ and prefrontal cortex.^55^ These inputs can activate inhibitory networks that desynchronize local network activity, and modulating their strength could produce the correlation effects observed in the current study. Given the diversity of these intracortical signals, it might seem surprising that the arousal-related changes reported here should occur in such a low-dimensional space. Further investigation with selective control of feedback from different cortical areas will determine if, in fact, the impact of signals from these different cortical areas can be dissociated in A1.

## Materials and Methods

### Surgical procedure

All procedures were performed in accordance with the Oregon Health and Science University Institutional Animal Care and Use Committee (IACUC) and conform to standards of the Association for Assessment and Accreditation of Laboratory Animal Care (AAALAC). The surgical approach was similar to that described previously.^2, 9, 56, 57^ Five young adult male ferrets were acquired from an animal supplier (Marshall Farms). Head-post implantation surgeries were then performed in order to permit head-fixation during neurophysiology recordings. Two stainless steel head-posts were fixed to the animal along the midline using UV-cured dental composite (Charisma) or bone cement (Palacos), which bonded to the skull and to stainless steel screws that were inserted into the skull. After a two-week recovery period, animals were habituated to a head-fixed posture and auditory stimulation. At this point, a small (0.5 - 1 mm) craniotomy was opened above primary auditory cortex (A1) for neurophysiological recordings.

### Acoustic stimuli

Digital acoustic signals were transformed to analog (National Instruments), amplified (Crown), and delivered through a free-field speaker (Manger) placed 80 cm from the animal’s head and 30°contralateral to the the hemisphere in which neural activity was recorded (Figure 1). Stimulation was controlled using custom MATLAB software (https://bitbucket.org/lbhb/baphy), and all experiments took place inside a custom double-walled sound-isolating chamber (Professional Model, Gretch-Ken).

Natural sounds stimuli were presented in four different configurations. Set 1 consisted of 93, 3-sec samples (2.5 sec ISI, *n* = 3 sites), set 2 consisted of 306, 4-sec samples (1 sec ISI, *n* = 14 sites), set 3 consisted of 306, 1-sec samples (0.5 sec ISI, *n* = 6 sites), and set 4 consisted of 2, 3-sec samples of ferret vocalizations (2.5 sec ISI, *n* = 2 sites). In sets 1-3, the stimulus sets contained species conspecific and heterospecific vocalizations, speech, music, and environmental sounds chosen to sample diverse spectro-temporal statistics.^35^ All stimuli were presented at 65 dB SPL. During every experimental session, a subset of samples were repeated at least ten times (set 1: 3 samples, set 2: 18 samples, set 3: 18 samples, set 4: all samples), while the remainder were played only once. In order to study the trial-to-trial variability in neural responses, only the high-repeat sounds were included in this study. The order in which stimuli were presented was generated pseudo-randomly. Stimuli were played continuously until all sound samples in the library had been presented. In the case of set 1, the entire stimulus set was repeated 2-3 times. This meant that experiments lasted approximately 40 minutes. The full sound library can be accessed at https://bitbucket.org/lbhb/baphy). Some of data used in this study has been published previously.^2, 58^

### Neurophysiology

Upon opening a craniotomy, 1 - 4 tungsten micro-electrodes (FHC, 1-5 MΩ) were inserted to characterize the tuning and response latency of the region of cortex. Sites were identified as A1 by characteristic short latency responses, frequency selectivity, and tonotopic gradients across multiple penetrations.^59, 60^ Subsequent penetrations were made with a single (64-channel) or dual shank (128-channel) silicon electrode array.^34^ Electrode contacts were spaced 20 μm horizontally and 25 μm vertically, collectively spanning 1.05 mm of cortex. On each consecutive recording day, we changed the location of the electrode penetration to access fresh cortical tissue, expanding the craniotomy as necessary. Data were amplified (RHD 128-channel headstage, Intan Technologies), digitized at 30 KHz (Open Ephys^61^) and saved to disk for further analysis.

Spikes were sorted offline using Kilosort^62^ or Kilosort2 (https://github.com/MouseLand/Kilosort2). Spike sorting results were manually curated in phy (https://github.com/cortex-lab/phy). For all sorted and curated spike clusters, a contamination percentage was computed by measuring the cluster isolation in feature space. All sorted units with contamination percentage less than or equal to 5 percent were classified as single-unit activity. All other stable units that did not meet this isolation criterion were labeled as multi-unit activity.

### Pupillometry

During neurophysiological recordings, video of the ipsilateral pupil (relative to the recording hemisphere) was collected using an open source camera (Adafruit TTL Serial Camera) fitted with a lens (M12 Lenses PT-2514BMP 25.0 mm) whose focal length allowed placement of camera 10 cm from the eye. Contrast was increased using infrared illumination. Ambient light levels were fixed for each experiment at roughly 1500 lux to provide maximum dynamic range of pupil size.^2^ Pupil size was measured offline by fitting an ellipse to each video frame using using a custom machine learning algorithm (Python and Tensorflow). The minor axis of the fit ellipse was extracted and saved for analysis with neurophysiological data. Blinks were detected and excluded as in^2^ and pupil data was shifted by 750 ms relative to spike times in order to account for the lagged relationship between changes in pupil size and neural activity in auditory cortex and to allow for comparison with previous research.^31^

The pupil tracking algorithm itself utilized a deep learning approach. Our model architecture was based on DenseNet201,^63^ which is available through Keras (https://keras.io/). In order to transform the output of the model to pupil ellipse predictions, we added a single global pooling layer and a final prediction layer in which five pupil ellipse parameters (x-position, y-position, minor axis, major axis, and rotation) were fit to each video frame. In order to initialize model weights, the model was pre-trained on ImageNet,^64^ then fine-tuned using roughly 500 previously analyzed, nonconsecutive frames from video of the pupil of multiple different ferrets (data from^2^). Qualitatively, after this first round of training the model performed well, even on novel video frames of pupil from new animals. However, in cases where the pupil video quality was poor, or differed substantially from the video frames in our training data set, we noticed failures in the model predictions. To further improve the model, we employed an active learning procedure. For each new analyzed video, pupil ellipse fits were analyzed qualitatively by experimenters. If the fit quality was deemed poor, predictions for these frames were manually corrected and added to the training data set. The model was then retrained and the analysis rerun. The network became robust to varying levels of video quality and performed consistently without the need for user intervention. The code for this analysis is available at https://github.com/LBHB/nems_db.

Because arousal was not explicitly controlled, identical arousal states were not sampled from day to day, between different animals, or between different stimulus pairs within the same experimental session. To control for this variability, we created a normalized metric of pupil variance for each stimulus, and then computed the mean of this variance metric for each recording session. For each stimulus pair, trials were split in half based on median pupil size across all trials. Next, the difference of mean large pupil size and mean small pupil size was normalized by the standard deviation of pupil across the entire experiment. Therefore, for a stimulus pair that sampled a large range of pupil states (relative to pupil during that particular recording session) this metric was high. Finally, we averaged across all stimulus pairs at a recording site and compared this mean across recording sessions (Figure S7).

### Pupil-dependent GLM

In order to characterize the dependence of first-order response statistics on pupil-indexed arousal, we built a state-dependent generalized linear model of sound-evoked activity. For each recorded unit, *i*, the input to this model was defined as the peri-stimulus time histogram (PSTH) response averaged over all stimulus repetitions (*r*_0*,i*_(*t*)). The predicted firing rate was calculated by scaling the PSTH by a pupil-dependent multiplicative and additive factor to model pupil-dependent changes in gain and baseline firing rate over time (Eqn. 1). To account for a possible nonlinear relationship between pupil size and neuromodulation, the pupil signal was first passed through a static sigmoid nonlinearity, *F* (double exponential^65^). The parameters of this nonlinearity, as well as the gain (*β*_0_) and baseline (*β*_1_) coefficients were fit independently for each cell using 10-fold jackknifed cross validation using the Neural Encoding Model System (NEMS, https://github.com/LBHB/NEMS).

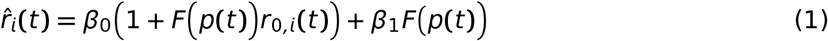

The sigmoid transformation applied to the pupil improved model performance, but made it difficult to interpret the gain (*β*_0_) and baseline (*β*_1_) parameters directly. To assess the relative magnitude of pupil effects on baseline and gain, we computed the unique modulation of firing rate due to each, respectively, as the mean difference in prediction between large and small pupil conditions:

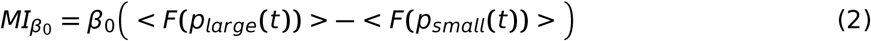

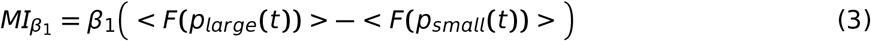

where *<>* indicates the mean over time and *large* / *small* refer to median splits of pupil size across the experimental session.

To quantify overall pupil-dependent modulation without differentiating between baseline and gain, we measured an overall pupil modulation index (*MI*). *MI* was defined by as the mean sound-evoked response when pupil was large minus the mean response when pupil was small, normalized by the sum of these two quantities. Large and small trials were defined based on a median split of pupil size across the entire recording session.

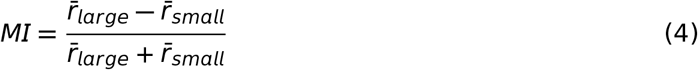

### Noise correlations

Pairwise noise correlations were measured by grouping spike counts into 250 ms bins, extracting only evoked periods (epochs when sound stimuli were playing), and computing Pearson’s correlation between all pairwise combinations of z-scored spike counts. Z-scores were calculated for each stimulus independently, as in Eqn. 5, where *r*(*t*) is the single trial response, *r*_0_ is the trial averaged response, and *σ* is the standard deviation of spike counts across repetitions. Therefore, the z-scored spike counts *Z*(*t*) of each neuron *i* for each stimulus *s* had mean zero and standard deviation one.

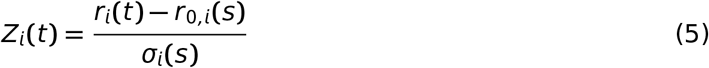

Noise correlations were also measured after first band-pass filtering spike counts. In order to achieve high frequency resolution, data were binned at 100 Hz (10 ms bins) for this analysis. To band-pass filter the data, we first computed the Fast Fourier transform of each unit’s spike counts. Next, in the frequency domain we used a Tukey window to extract the frequencies of interest. Finally, we inverted the modified signal back into the time domain, z-scored the data, and computed Pearson’s correlation as above. For pupil-corrected noise correlations, we performed the same procedure after first performing the pupil-correction (see below).

### Pupil-correction

To remove changes in firing rate that were associated with pupil, we regressed out all stimulus-independent variability that could be explained with pupil size. We performed this correction on a per-neuron, per-stimulus basis to completely remove pupil-associated changes in firing.

For each neuron (*i*) and stimulus (*s*) the mean response across all trials was measured 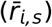 and subtracted from the true response (*r_i,s_*, Eqn. 6). A linear regression model was used to predict to the residual variability in firing rate with pupil size (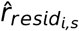, Eqn. 7). The prediction of this model was then subtracted, and the remaining, pupil-corrected, trial to trial variability was added back to the mean stimulus response (Eqn. 8).

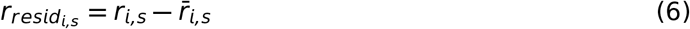

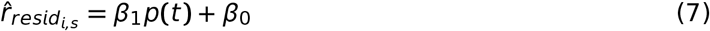

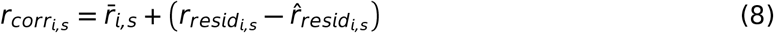

### Pairwise stimulus discrimination

Natural sound samples were broken into non-overlapping 250 ms segments, similar to the procedure followed by Pachitariu et al., 2015.^66^ In order to obtain cross-validated estimates of stimulus discriminability (Figure S2), we required that each stimulus be repeated at least 20 times. This was the case for 11 / 25 recording sites. These 11 recording sites were collected from four different animals. For the remaining sites (which contained only 10 repetitions per stimulus), we could not perform cross-validation. Therefore, for all analyses included in the main text in which we report neural discriminability (*d*′^2^) estimates (Figures 2, 3, 4, and 6) only the high-repetition count recording sessions were included. However, we repeated key analyses using all 25 recording sessions without performing cross-validation and found that results were largely consistent with the more rigorous approach (Figure S8).

For each pair of stimulus segments we extracted the *N* neuron X *k* trial response matrices, *A* and *B*. Because the number of recorded neurons was greater than the number of stimulus repetitions, we performed *dDR* to preserve only significant dimensions of the population response (see following section and Figures S3/S4). This allowed us to accurately estimate the population statistics. We quantified encoding accuracy in this reduced-dimensionality space by measuring neural stimulus discriminability, *d*′^2^, the discrete analog of Fisher information:^26, 27, 32, 37–39^

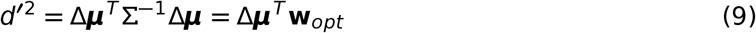

where Δ***μ*** represents the vector connecting the mean ensemble responses to stimulus A and stimulus B, 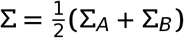 represents the mean noise-covariance matrix, and **w**_*opt*_ is the optimal decoding axis, *i.e.* the vector orthogonal to the optimal linear discrimination hyperplane in state-space. In practice, for our high repetition count data set we estimated **w**_*opt*_ using 50-percent of trials (training data) then projected the held out 50-percent of trials (test data) onto this vector and measured discriminability. For a detailed schematic of this procedure, see Figure S2. Pupil-dependent measurements of *d*′^2^ followed an identical procedure, but before measuring discriminability, the test data was first split in half based on median pupil size.

Throughout this work, we describe stimulus discriminability as a function of the relationship between the signal and noise subspace. To quantify this relationship, we defined the following two axes which we found to be critical for understanding encoding accuracy:

#### Discrimination axis magnitude (|wΔμ|)

For a given pair of stimuli A and B, we defined the discrimination axis (*i.e.* the axis containing sensory information) to be the vector Δ***μ***. When two stimuli drove very different population activity, the magnitude of this vector, ***|***Δ***μ|***, was large and discriminability was high.

#### Noise interference (| cos(θ_Δμ,e_1__)|)

The structure of trial to trial variability in responses can vary depending on the stimulus.^25, 67, 68^ Therefore, within the reduced dimensionality *dDR* space (see below) for each pair of stimuli, we defined the correlated variability axis, **e**_1_, as the first eigenvector of the average covariance matrix, 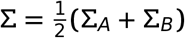. We then asked how this axis interacted with the signal by measuring the cosine similarity between Δ***μ*** and **e**_1_. Thus, when the two were perfectly aligned our measure of noise interference was equal to one and when they were perfectly orthogonal, interference was zero.

To quantify the dependence of *d*′^2^ and m*d*′^2^ on these two axes, we used the following linear regression models (Eqns. 10, 11). All variables were z-scored prior to model fits.

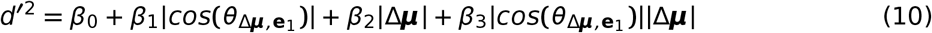

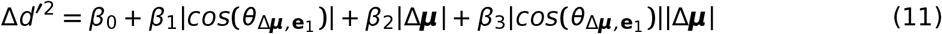

### Decoding-based Dimensionality Reduction

During our experiments we typically recorded from between 20 and 60 units simultaneously at a single recording site (on one 64-channel probe shank). However, at most, we repeated each individual stimulus only 24 times. Therefore, because the number of neurons was generally greater than the number of trials, estimation of the covariance matrix for a particular stimulus, A (∑_*A*_), was unreliable (Figure S4).^32^ This poses a challenge for estimating neural discriminability as this process depends on accurate estimation of ∑. One technique that has been proposed to help deal with such limitations is dimensionality reduction.^32^

Because we were interested specifically in studying neural discriminability, we developed a decoding-based Dimensionality Reduction (*dDR*) approach whereby we reduced our *N*-dimensional space, on a per-stimulus pair basis, to just two dimensions containing the most critical information for estimating stimulus discriminability: The sensory discrimination axis, Δ***μ***, and the global noise axis, which we defined as the first principal component of the pooled noise data over all stimuli and neurons (raw spike counts after subtracting the mean stimulus response from each neuron).^23^ Thus, the noise axis was fixed between stimulus pairs but Δ***μ*** was not. We defined *dDR*_1_ = Δ***μ*** and *dDR*_2_ as the axis orthogonal to Δ***μ*** and in the plane spanned by Δ***μ*** and the noise axis. We illustrate this process graphically (Figure S3). Analysis of neural discriminability using *dDR* in simulated data sets (Figure S4) demonstrated the efficacy of this method both in preventing over-fitting and in facilitating accurate estimation of discriminability. Furthermore, though it is not a focus of the current work, we demonstrate that although this method only explicitly preserves the first noise principal component, it is still capable of detecting information limiting correlations^26^ that recent work has suggested might be restricted to relatively low variance dimensions of the data.^32, 38^

### Pupil-dependent changes in stimulus discriminability

*d*′^2^, measured across pupil states, could vary greatly across the sensory response. Therefore, in order to measure pupil-dependent changes in coding accuracy, we used a normalized metric, m*d*′^2^ that allowed us to directly compare relative changes in discriminability for different stimulus pairs and recording sites. To measure this, we computed a pupil-dependent modulation index of *d*′^2^. For each stimulus pair, m*d*′^2^ was defined as the *d*′^2^ measured during large pupil trials minus *d*′^2^ for small pupil trials, normalized by the sum of these two quantities.

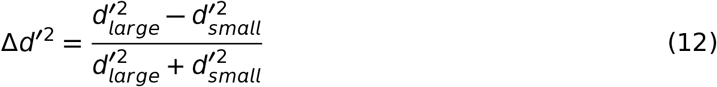

### Contribution of single neuron and correlated variability to changes in discriminability

We observed that pupil modulated both the mean distance between stimuli (discrimination axis / signal magnitude) and the amount of trial to trial variability in the response to the same stimulus. To determine which contributed to pupil-dependent changes in *d*′^2^, we performed a linear regression between m*d*′^2^ and the pupil-dependent changes in both signal magnitude and shared noise variance (Eqn. 13). We also included an interaction term to account for changes in the alignment of the noise and signal axes between arousal states. Each of these statistics was calculated on a per stimulus-pair basis, and the models were fit per experimental session to prevent bias arising from any differences between experiments.

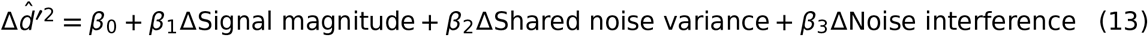

In order to determine if each individual variable explained significant variability in m*d*′^2^, we created three different single variable regression models, one for each of the population statistics in the full model (Eqn. 13). We then measured cross-validated estimates of *R*^2^ for each (*cvR*^2^). To do this, models were fit on 90-percent of the data and evaluated on the remaining 10-percent. We used jackknifing so that cross-validated predictions tiled the entire dataset. A single *cvR*^2^ value was measured for each experiment using this prediction. Finally, to determine if the effect of a given predictor was consistent across experiments, we tested whether the distribution of *cvR*^2^ values across experiments was significantly different than zero.

### Response simulations

To dissociate arousal-dependent modulation of single neuron response statistics, such as response gain, from correlated variability effects, we followed a procedure similar to that in Cohen et al., 2009.^16^ Simulated data sets were generated independently for both large and small pupil conditions. In the independent variability simulation, the mean and variance of single neurons were allowed to vary between large and small pupil while correlations between neurons were fixed. In the full simulation, correlations between neurons were also allowed to change between large and small pupil.

#### Independent variability simulation

Pupil-dependent mean and variance of single neurons were measured using the raw data. Next, we built covariance matrices, ∑_*ind., large*_ and ∑_*ind.,small*_, where the off-diagonal elements in each pupil condition were fixed over all pupil conditions and the diagonal elements of each matrix were set to the measured variance in each respective pupil condition. We simulated population responses, *R_ind_* by drawing trials from a multivariate Gaussian distributions as follows:

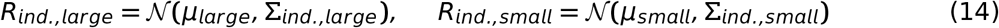

#### Full simulations

Pupil-dependent single neuron statistics (mean and variance) *and* correlated variability between neurons (off-diagonal elements of the covariance matrix) were measured using the raw data. We used these values to define covariance matrices Δ_*full,large*_ and ∑_*full,small*_ where all elements of the matrices were set based on their covariance values in each respective pupil condition. We simulated full population responses, *R_full_* as above by drawing trials from a multivariate Gaussian distributions. All simulations were performed on a per-stimulus basis.

### Identification of noise correlation modulation axis

We identified the axis in state space along which noise correlations were most modulated by pupil. First, we measured large and small pupil noise covariance matrices. Noise covariance matrices were defined as the covariance matrix of the z-scored spiking activity (z-scored on a per-stimulus basis). We then set the diagonal of this matrix equal to zero, because for this analysis we were interested only in capturing shared patterns of variability. Finally, we computed the difference covariance matrix (small pupil matrix minus large pupil matrix) and performed an eigen-decomposition. The largest eigenvalue corresponded to the axis along which noise correlations changed most between the two pupil states. Significance of this axis was evaluated with a permutation test.

### Statistical analyses

Our data followed a nested structure; multiple cells were recorded from the same animal and many different stimuli were presented during each experimental session. Therefore, it is possible our results could be biased by differences between animals and/or experimental recording session. To account for this, in all of our statistical tests we took one of the following two approaches: (1) Averaged metrics across cells (or pairs of cells) and sound stimuli within a recording session before performing statistical tests or (2) Performed statistical tests using hierarchical bootstrapping.^69^ Although each approach reduces statistical power relative to treating each individual measurement as wholly independent, they provide conservative estimates of p-values and reduce the chance of detecting false positives.^69^ The only exception to this approach was in the estimates of pupil-dependent effects on evoked firing rates in single neurons shown in Figure S1.

For estimating significance of pupil-effects in single neurons, we performed a jackknifed t-test for each individual neuron. Data were split into 20 non-overlapping estimation / validation sets that together tiled the entire experimental session. For each validation set, a prediction correlation was computed for both the full pupil-dependent GLM and the pupil-shuffled GLM. Cells where the mean pupil-dependent model performance was 2 standard errors greater than the mean shuffled model performance (*p **≤*** 0.05) were considered to have significant pupil effects.

For all statistical tests measuring large vs. small pupil effects where we first averaged results within recording session, we performed a two-tailed Wilcoxon signed-rank test. For each test, we report the test statistic, *W*, the p-value, and the exact *n* number of recording sessions used to perform the test. In cases where we performed a hierarchical bootstrap, we report the direct bootstrap probability of the null hypothesis.^69^ In both cases, we also provide the mean and standard error of the number of measurements per recording session

To quantify patterns in *d*′^2^ and m*d*′^2^ heatmaps, we performed standard linear regression on a per-recording site basis using the statsmodel package for Python (https://www.statsmodels.org/stable/index.html). To determine the significance of regression coefficients at the group level, we performed a Mann-Whitney U test to determine if the parameter distribution over recording sites was significantly different than zero. For each test, we report the test statistic, *U*, the p-value, and the exact *n* number of recording sessions used to perform the test.

To measure the significance of the correlation between pupil-dependent changes in stimulus discriminability and pupil variance per recording session, we performed a permutation test. We randomly shuffled our distribution of mean m*d*′^2^ across recording sites and computed the correlation of these re-sampled values with pupil variance. This was repeated 1000 times in order to calculate a p-value, which is reported in the text along with the true measured correlation coefficient.

Finally, to determine if the measured noise correlation modulation axis for a given recording session was significant, we performed a permutation test. For each recording session, pupil size was shuffled in time before classifying trials as large or small pupil. After shuffling, spike covariance matrices were measured for large and small pupil trials and an eigen-decomposition was performed on the difference between these matrices. This was repeated 20 times and the mean / standard error of the resulting eigenvalues was stored. Significant dimensions were those in which the actual measured eigenvalue was 2 standard errors greater than the mean shuffled value.

## Glossary

Discrimination/signal axis (Δ**μ**): Vector connecting the mean population response to stimulus A and stimulus B.
Pooled noise data: All single trial data after subtracting off mean stimulus responses from each neuron.
Global noise axis: First principal component of pooled noise data.
Correlated variability axis (**e**_1_): Stimulus-pair specific estimate of noise axis. First principal component of noise data pooled across the two stimuli within dDR space.
Optimal decoding axis (**w**_opt_): Axis orthogonal to the optimal linear discrimination boundary in dDR space.
Discrimination axis magnitude (**|**Δ**μ|**): Vector magnitude of the discrimination axis.
Noise interference (**|** cos(θ_**e**1_,Δ**μ**)|): Cosine similarity between the correlated variability axis and discrimination axis.

## Data Availability

The datasets analyzed in this study are available from the corresponding author upon reasonable request.

## Author contributions

CRH and SVD designed experiments. CRH performed experiments, analyzed data, and developed pupil tracking software. DS assisted with data collection. ZPS assisted with software development for pupil tracking and assisted with data collection. CRH and SVD wrote the manuscript. All authors edited the manuscript.

## Acknowledgments

This work was supported by a National Science Foundation Graduate Research Fellowship (NSF GRFP, GVPRS0015A2) (CRH), the National Institute of Health (NIH, R01 DC0495) (SVD), Achievement Rewards for College Scientists (ARCS) Portland chapter (CRH), and by the Tartar Trust at Oregon Health and Science University (CRH).

