## Supplementary Material for "Selective effects of arousal on population coding of natural sounds in auditory cortex"

*Heller et al.*

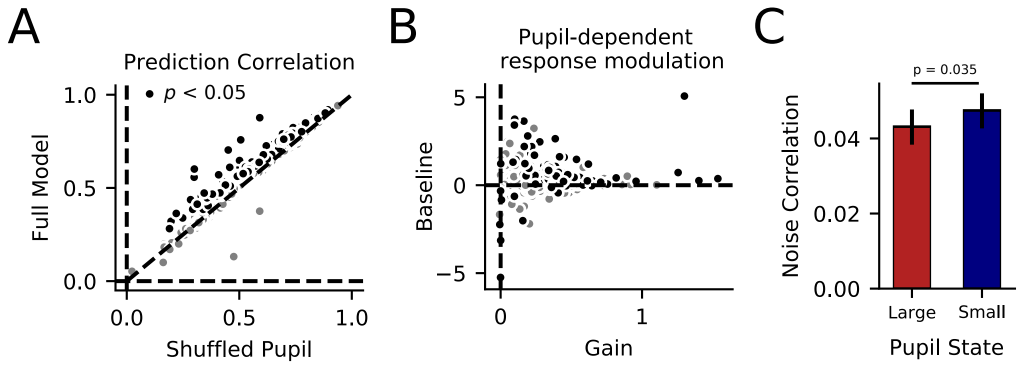


**Figure S1. Pupil-indexed arousal increases response gain and decreases pairwise noise correlation in A1. A.** Performance of a pupil-dependent GLM (Eqn. 1) was evaluated by computing prediction correlation (Pearson's $r$) between predicted and the actual firing rate. For 105/702 recorded units, the full model (pupil-dependent GLM) significantly outperformed the sensory-only model, where pupil size was shuffled in time (Jackknifed t-test, $p < 0.05$). Most other neurons trended toward an improvement, and the effect was significant across the population (full model mean $r = 0.65 \pm 0.02$, shuffled pupil $r = 0.63 \pm 0.02$, $p \approx 0.0$, Bootstrap test, $n = 25$ recording sessions, $31.9 \pm 5.17$ units per session). **B.** Mean change in gain or baseline between large and small pupil (Eqns. 2, 3). Both values were consistently greater than zero (gain modulation: $\mu = 0.21 \pm 0.02, p \approx 0.0$, baseline modulation: $\mu = 0.24 \pm 0.05, p \approx 0.0$, Bootstrap test, $n = 25$ recording sessions, $31.9 \pm 5.17$ units per session). **D.** Mean noise correlation for all simultaneously recorded pairs of units, grouped within recording session and averaged across sessions ($n = 25$ recording sessions, $318 \pm 94$ unit-pairs per session), computed over trials when pupil was large versus trials when pupil was small. Error bars represent mean / standard error across recording sessions. Noise correlation was significantly reduced during large pupil trials ($\rho_{small} = 0.047 \pm 0.005$ vs. $\rho_{large} = 0.043 \pm 0.005$, $p = 0.035$, Bootstrap test).

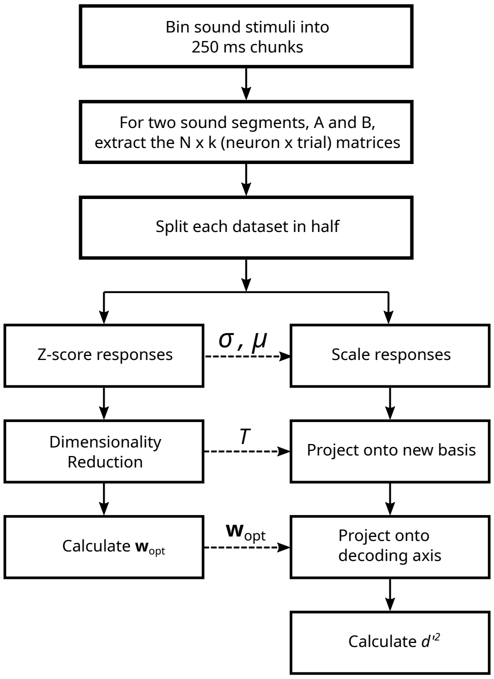


**Figure S2. Calculating neural discriminability.** Procedure for measuring discriminability of natural sound segments by neural populations in primary auditory cortex (A1). Stimuli were binned into 250 ms chunks, and a discrimination index, $d'^2$, was calculated between each pairwise combination of segments. To prevent over-fitting to noise in the data, we took two steps: First, we performed cross-validation, estimating the optimal decoding axis using 50% of the data and evaluating $d'^2$ along this axis with the held out 50%. Second, because we were in a trial-limited regime where the number of stimulus repetitions, $k$, was less than the number of neurons, $N$, we performed dimensionality reduction, reducing the number of effective dimensions from $N$ to two, allowing us to reliably estimate neural discriminability for each pair of stimuli (see Figure S4).

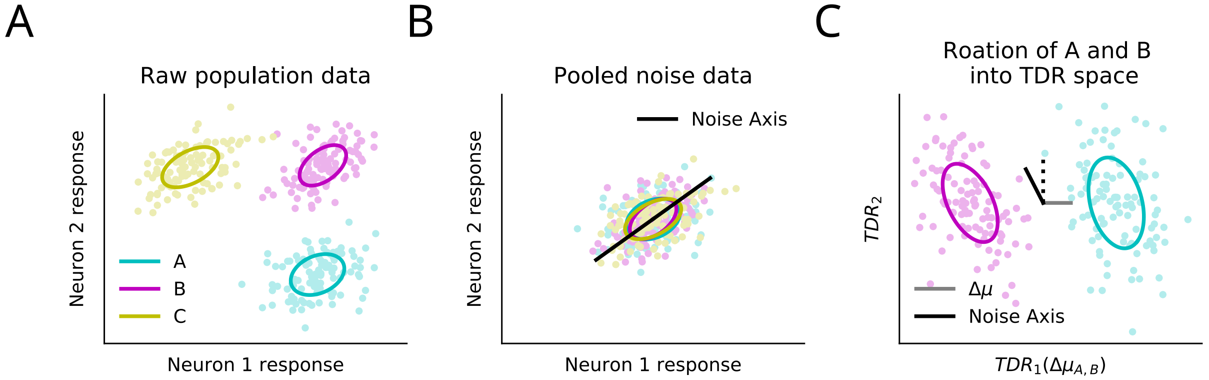


**Figure S3. decoding-based Dimensionality Reduction (dDR) procedure.** We illustrate a simple rotation from a two-dimensional neuron state space into *dDR* space, but the same logic applies for larger neural populations. **A.** Simulated responses to three stimuli (A, B, and C) for two neurons. Dots represent single-trial responses, and ellipses summarize the standard deviation of trial-trial variability in both neurons. **B.** Pooled noise data is defined by subtracting the mean response to each stimulus and pooling across all stimuli. Then, we define the noise axis as the first principle component of this pooled data. **C.** The *dDR* space for each pair of stimuli is defined by two dimensions: The difference in the mean response to two stimuli, $\Delta\mu$, and the axis orthogonal to $\Delta\mu$ in the plane defined by $\Delta\mu$ and the noise axis (dashed line / y-axis).

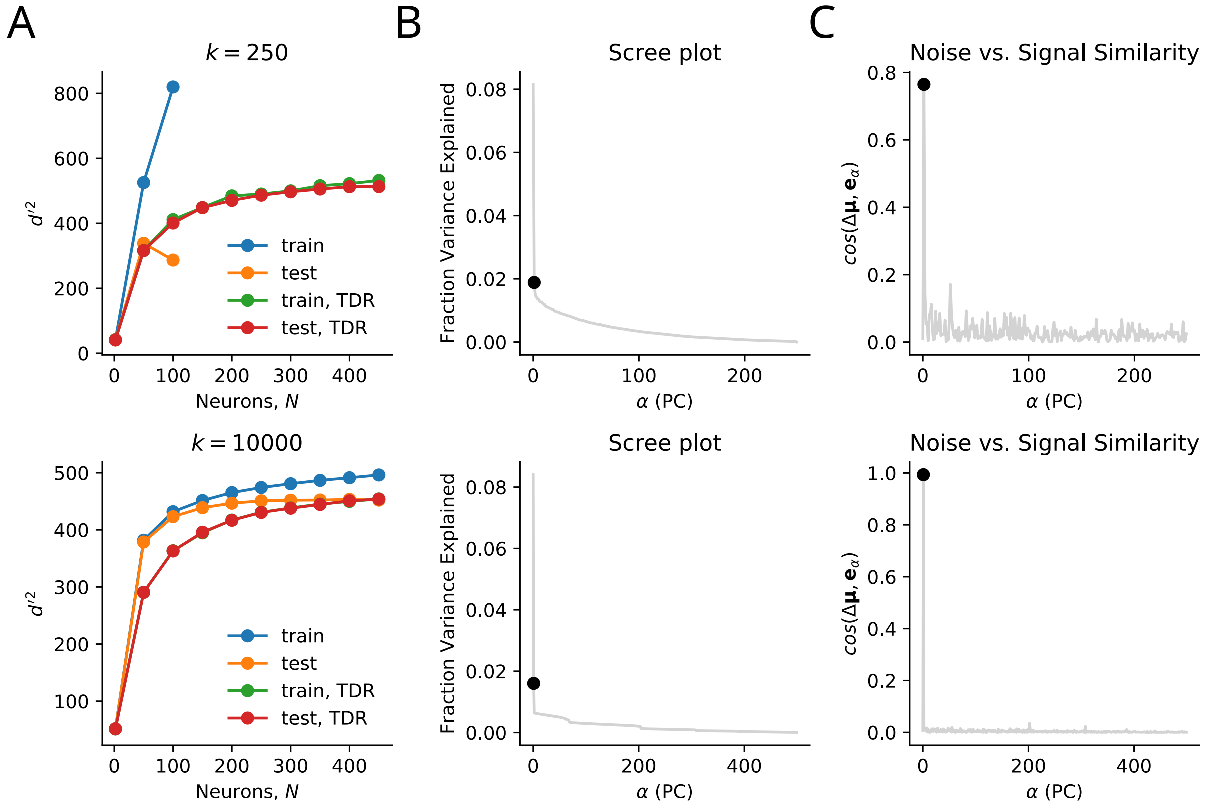


**Figure S4. Targeted Dimensionality Reduction prevents over-fitting and detects weak, information limiting correlations.** Two data sets were simulated. The first illustrates a trial limited regime where the number of trials, $k$, is less than the number of neurons, $N$ (top). The second illustrates the case where $k >> N$, and estimation of information using the whole data set is feasible. The simulations were constructed such that the covariance matrices had two significant principal noise components, one of which contained information limiting correlations. **A.** $d'^2$ is shown as a function of the number of neurons for the two simulated data sets: one with $k << N$ (top) and one with $k >> N$ (bottom). In the trial limited regime, calculating $d'^2$ using the full data set (without dimensionality reduction) leads to drastic over-fitting, with very high $d'^2$ in the training data (blue) but low $d'^2$ in the test data (orange). *dDR* prevents this problem, with a good match between training and test data. Furthermore, the *dDR* results show that information saturates as neurons are added, evidence of information limiting correlation. In the case of $k >> N$, when we have sufficient data to use the full population, *dDR* performance approaches the asymptotic value of information for the full population (orange line). This same performance is reached even in the trial limited regime (top). **B.** Scree plots show the fraction variance of population activity explained by each noise component. Black dot indicates the component aligned with the discrimination axis ($\Delta\mu$) *i.e.* containing information limiting noise. **C.** The alignment of each noise component with $\Delta\mu$. Black dot indicates the component containing information limiting noise. Even though this noise is not contained in the first principal component, *dDR* is still able to detect it.

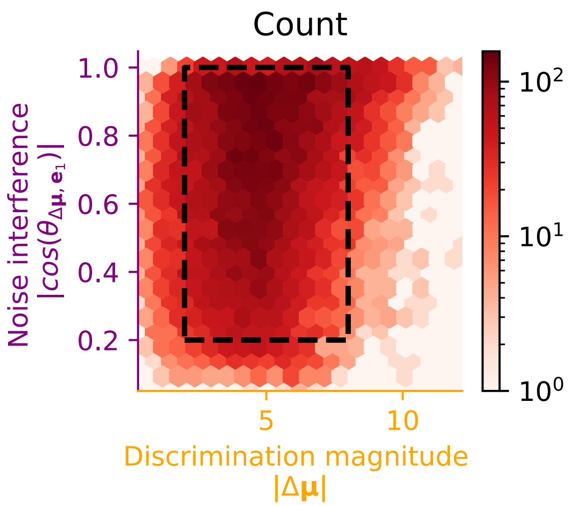


**Figure S5. Stimulus pairs span sensory vs. noise response space.** Histogram of natural sound pairs, binned according to their discrimination axis magnitude (x-axis) and their noise interference (y-axis). Color map is on a log scale. In figures 2, 3, and 4, $d'^2$ and $\Delta d'^2$ heatmaps were only plotted for bins in which every recording session had data present (black dashed box). Unless otherwise indicated, for all other plots and statistical tests all data were included.

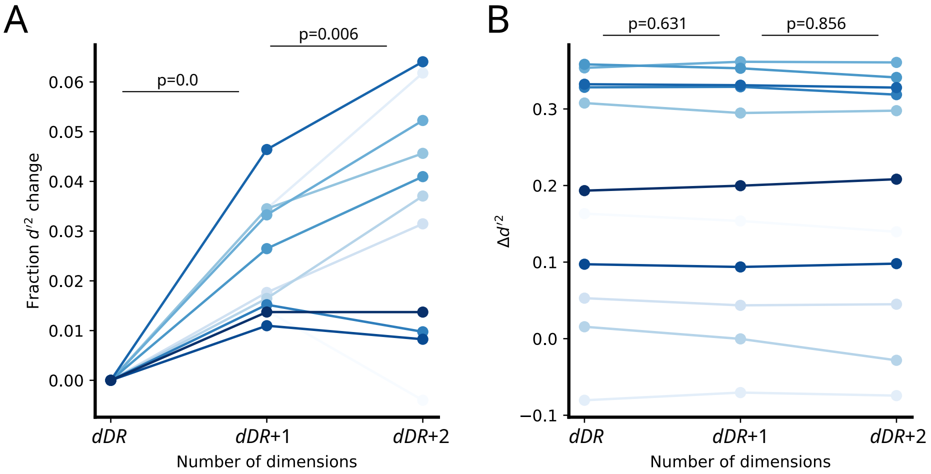


**Figure S6. Effect of dDR dimensionality on $d'^2$ estimates.** In addition to standard dDR (single noise dimension), $d'^2$ was computed for spaces with one or two additional noise dimensions. **A.** Each line indicates mean $d'^2$ for a single recording session, normalized to its value for the standard two-dimensional *dDR* case. Only high-repetition recording sessions, where cross-validation was possible, were included in this analysis (Methods). The percent-increase in overall $d'^2$ was significant for a single additional dimension (2.4 percent-increase, $p \approx 0.0$, Bootstrap test), and for adding a third noise dimension (0.8 percent-increase, $p = 0.006$, $n = 11$ recording sessions, Bootstrap test). However, for some individual sites, adding a third noise dimension led to over-fitting (reduced $d'^2$). **B.** Mean measurements of $\Delta d'^2$ for each recording session are shown as a function of the same three *dDR* spaces shown in (A). There is no significant difference between the standard two-dimensional *dDR* space, and *dDR* spaces with extra noise dimensions included ($n = 11$ recording sessions, Bootstrap test)

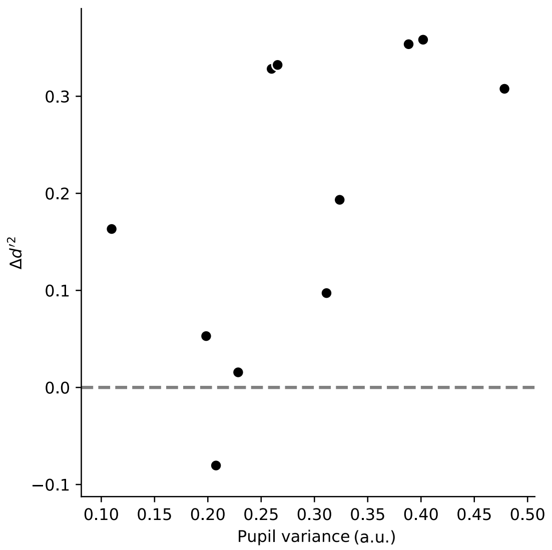


**Figure S7. Arousal-dependent improvements in coding accuracy scale with variability in pupil size.** Scatter plot compares $\Delta d'^2$ versus pupil size variability per recording site. These quantities were significantly correlated ($r = 0.63, p = 0.029, n = 11$ recording sessions, permutation test). Pupil size variability was measured for each 250-ms stimulus segment as the difference between mean large pupil and mean small pupil (split at the median) divided by the standard deviation of pupil across the entire experiment. Mean variability was then computed across all stimuli for each recording session.

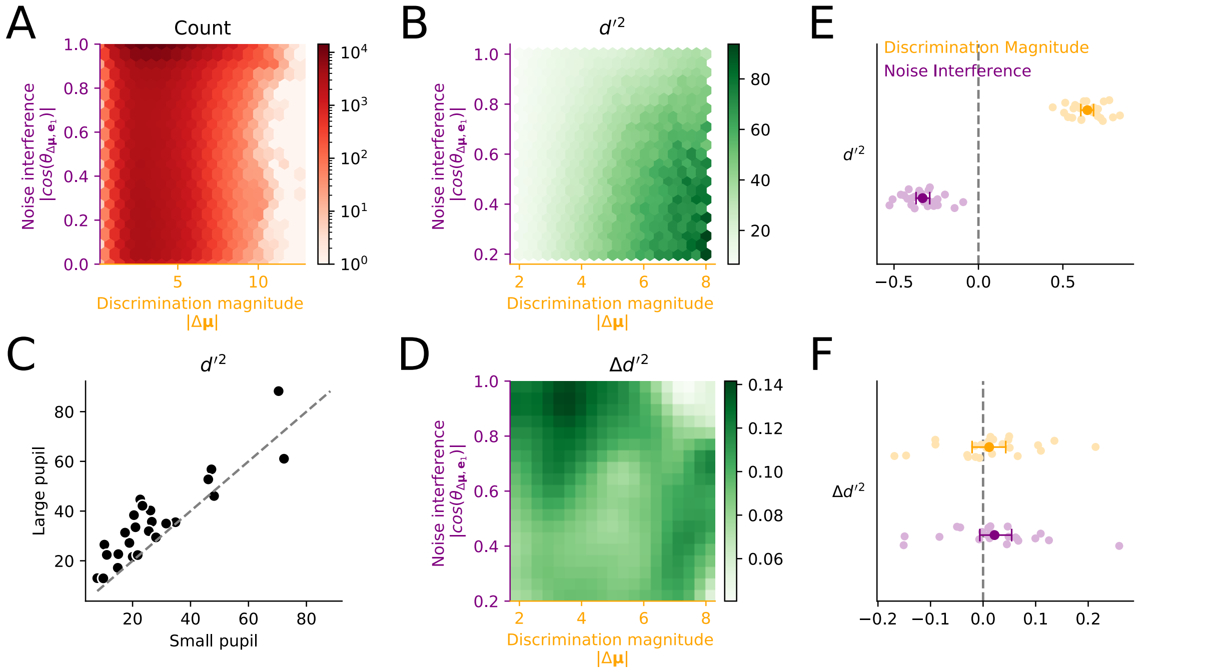


**Figure S8. Fit-data results follow similar patterns to cross-validated results.** For a subset of our experiments, we did not present each stimulus a sufficient number of times to perform rigorous cross-validation of $d'^2$ estimates. Although a less rigorous test, we re-performed all analysis from Figures 2 and 3 using all datasets and not performing cross-validation. **A.** Stimulus-pair count histogram, as in Figure S5. **B.** $d'^2$ binned and plotted as in Figure 2A. **C.** Mean large and small pupil $d'^2$ for each recording session as in Figure 3A. Large pupil $d'^2$ was significantly greater than small pupil ($p = 0.0001, W = 21, n = 25$ recording sessions, Wilcoxon signed-rank test). **D.** $\Delta d'^2$ binned and plotted as in Figure 3B. **E.** Regression coefficients predicting $d'^2$ for each recording session as in Figure 3E. Error bars represent the bootstrapped 95% confidence interval over recording sessions. $\beta_{Noise} = -0.33 \pm 0.02, p = 1.33e-9, U = -6.06, \beta_{Discrimination} = 0.65 \pm 0.02, p = 1.33e-9, U = 6.06, n = 25$ recording sessions, Mann-Whitney U test). **F.** Regression coefficients predicting $\Delta d'^2$ for each recording session as in Figure 3E. Error bars represent the bootstrapped 95% confidence interval over recording sessions. $\beta_{Noise} = 0.022 \pm 0.016, p = 0.007, U = 2.66, \beta_{Discrimination} = 0.011 \pm 0.016, p = 0.22, U = 1.21 n = 25$ recording sessions, Mann-Whitney U test).

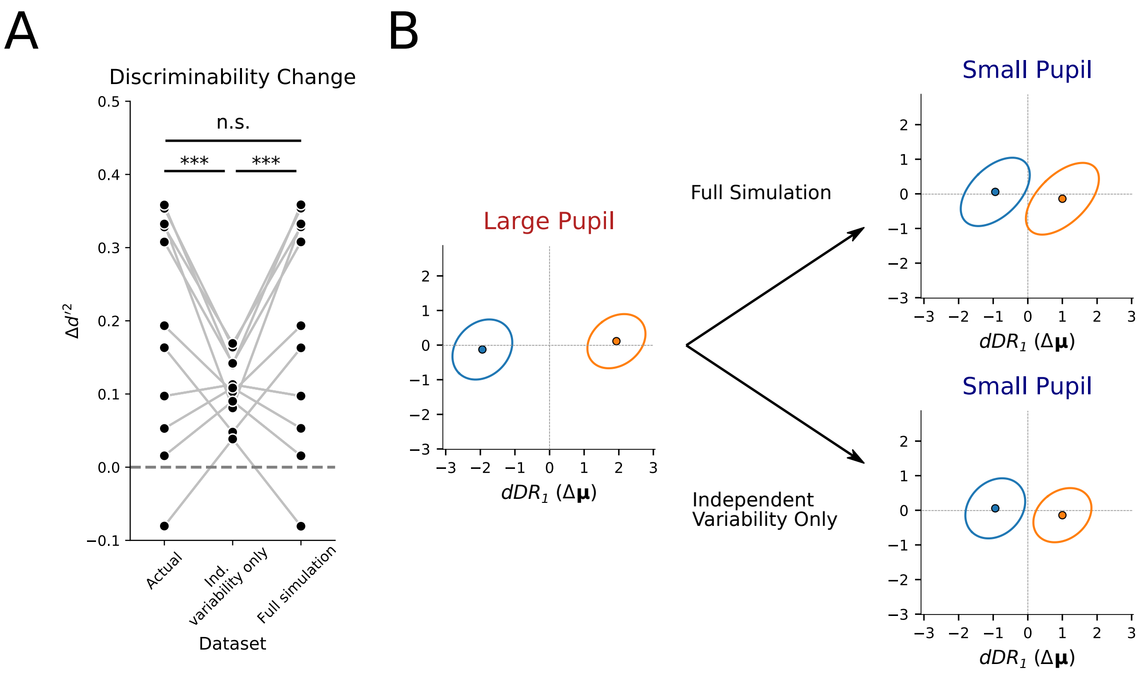


**Figure S9. Pupil-dependent changes in single neuron mean and variance, as well as correlated variability, account for changes in discriminability. A.** $\Delta d'^2$ for each recording session ($n = 11$) for the actual data (left), the independent variability only simulation (middle, Eqn. 14), and the full simulation (right). The full simulation, which included pupil-dependent changes in correlated variability, accurately reproduced the effects observed in the actual data ($p = 0.09$ Bootstrap, $n = 11$) while the independent variability only simulation did not ($p = 0.0002$ Bootstrap test, $n = 11$) **B.** Cartoon schematic of response distributions to two stimuli (orange vs. blue) in the *dDR* space, as in Figure 4. In the independent variability only simulation, correlated variability (ellipse shape) does not change between pupil conditions, but mean response (ellipse location) does.

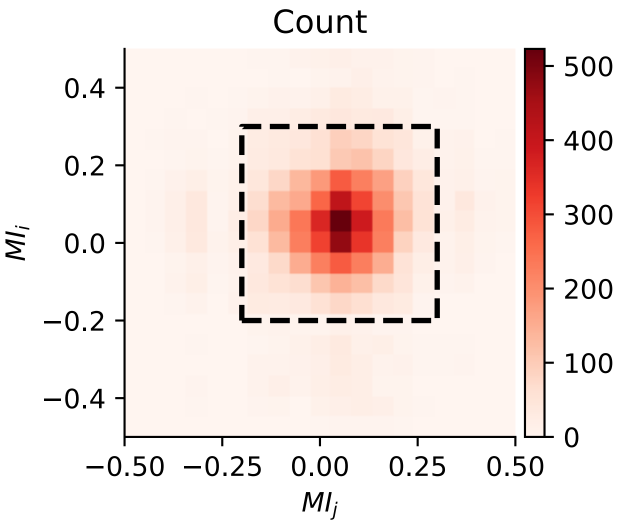


**Figure S10. Distribution of first-order pupil modulation indices for neuron pairs.** Histogram of simultaneously recorded unit pairs as a function of their first-order pupil modulation index, *MI* (Eqn. 4). Only pairs bounded by the black dashed box were included for the heatmap in Figure 5, however, all data was included for statistical tests.
